# Nautilus shell morphomics reveals microstructural heterogeneity alongside structural continuity across component boundaries

**DOI:** 10.64898/2026.07.31.742026

**Authors:** Kazuki Hirota, Takenori Sasaki, Davin H. E. Setiamarga

**Author notes:** Corresponding author (DHES).

## Abstract

The nautilus (*Nautilus* sp.) is an early-branching cephalopods. It retains several conchiferan synapomorphies, including an external planispiral biomineralized shell. The shell’s complex structure allows it to withstand hydrostatic pressure, control buoyancy, and protect against external hazards. In this study, we comprehensively examined shell microstructures across different shell components and regions representing various ontogenetic stages in two adult museum shell specimens. We found that the nautilus shell is composed of five microstructural types (spherulitic, prismatic, nacreous, semi-prismatic, and irregularly oriented prismatic structures) organized into layered architectures within individual shell components and coordinated across the shell as an integrated system. Our observations highlight transitions between distinct microstructures within and across shell components and local variation within individual components such as the dorsal and ventral shell walls, suggesting that these patterns may contribute to shell strength and overall mechanical performance. Variation in caecum morphology suggests that this structure may be developmentally plastic and subject to relatively relaxed structural constraints. These findings show that the *Nautilus* shell is an integrated biomineral system in which diverse microstructures are organized across shell components to meet functional demands and provide the mechanical strength needed for survival.

## Introduction

Nautiloidea is an early-branching cephalopod subclass that originated more than 400 million years ago in the Paleozoic (Walker and Brett 2002). Global nautiloid fossil records document an extensive diversification of shell morphology, including straight, curved, and coiled forms (Evans et al. 2014). Fossil evidence also indicates that nautiloids were already diverse in the Ordovician and dominated benthic–pelagic marine ecosystems through much of the Paleozoic (Crick 1981; Frey 1995). Their diversity began to decline after the Devonian and was severely reduced during the Permian–Triassic mass extinction, when many Paleozoic lineages disappeared, and although several groups persisted into the Triassic, their diversity continued to decline throughout the Mesozoic (Kummel 1953; Kummel 1956; Kröger et al. 2011; Kröger 2013; Combosch et al. 2017). A small number of lineages persisted into the later Mesozoic and Cenozoic, ultimately leaving only the extant lineage (Teichert and Matsumoto 2010). Studies have suggested that the nautilus and its extant relatives survived by occupying deep reef to continental slope habitats that buffered them from shallow-water extinction events (Saunders 1984a; Kiel et al. 2022).

Extant Nautiloidea represent the only surviving branch of this ancient cephalopod radiation and the only cephalopod lineage whose living members still retain a calcified external shell (Combosch et al. 2017; Ward et al. 2016; Setiamarga et al. 2021a, b; Hirota et al. 2023). The few species that persist today, such as the chambered nautilus (*Nautilus pompilius*), inhabit the tropical southwestern Pacific and the eastern Indian Ocean, mainly along continental slopes and outer reef environments at depths of approximately 100 to 700 m (Dunstan et al. 2011a; Barord et al. 2014; Barord et al. 2023). Morphological conservatism in the nautilus is evident in the retention of several ancestral features: a fully external shell, circumoral appendages lacking suckers, two pairs of gills, and the complete absence of an ink sac (Vermeij 1988; Shigeno et al. 2008; Sasaki et al. 2010). The animals rely primarily on chemosensory cues for foraging (Basil et al. 2000; Westermann and Beuerlein 2005; Basil et al. 2005; Barord et al. 2021) and show diel vertical migrations controlled by hydrostatic adjustments (Dunstan et al. 2011a). Their physiology is characterized by slow metabolism and low aerobic scope, consistent with a low-energy lifestyle in mesophotic and upper bathyal environments (Boutilier et al. 2000; Tajika et al. 2023). Buoyancy is regulated through the phragmocone chambers and siphuncular fluid exchange, a system that compensates for the absence of muscular fins found in coleoids (Denton and Gilpin-Brown 1966; Ward and Martin 1978; Greenwald et al. 1980; Greenwald et al. 1982).

The nautilus shell is formed of three major components, the shell wall, septa, and siphuncle, which together produce a chambered phragmocone supporting neutral buoyancy, structural strength, and mechanical stability (Lemanis et al. 2016; Karp et al. 2023; Fig. 1; Table 1). The siphuncle traverses each septum, allowing osmotic exchange of gases and fluids between chambers, which is mediated by secretory epithelium and connecting rings (Ward et al. 1981; Greenwald and Ward 2010). The geometric pattern of septal curvature and the logarithmic coiling of the shell confer resistance to hydrostatic pressure, allowing survival at depths exceeding 500 m (Westermann 1973; Kanie et al. 1980; Lemanis et al. 2016; Karp et al. 2023). This architecture is unique among living mollusks and represents a structurally complex functional design adapted for buoyancy control and hydrostatic pressure resistance (Lowenstam et al. 1984).

**Fig. 1.**
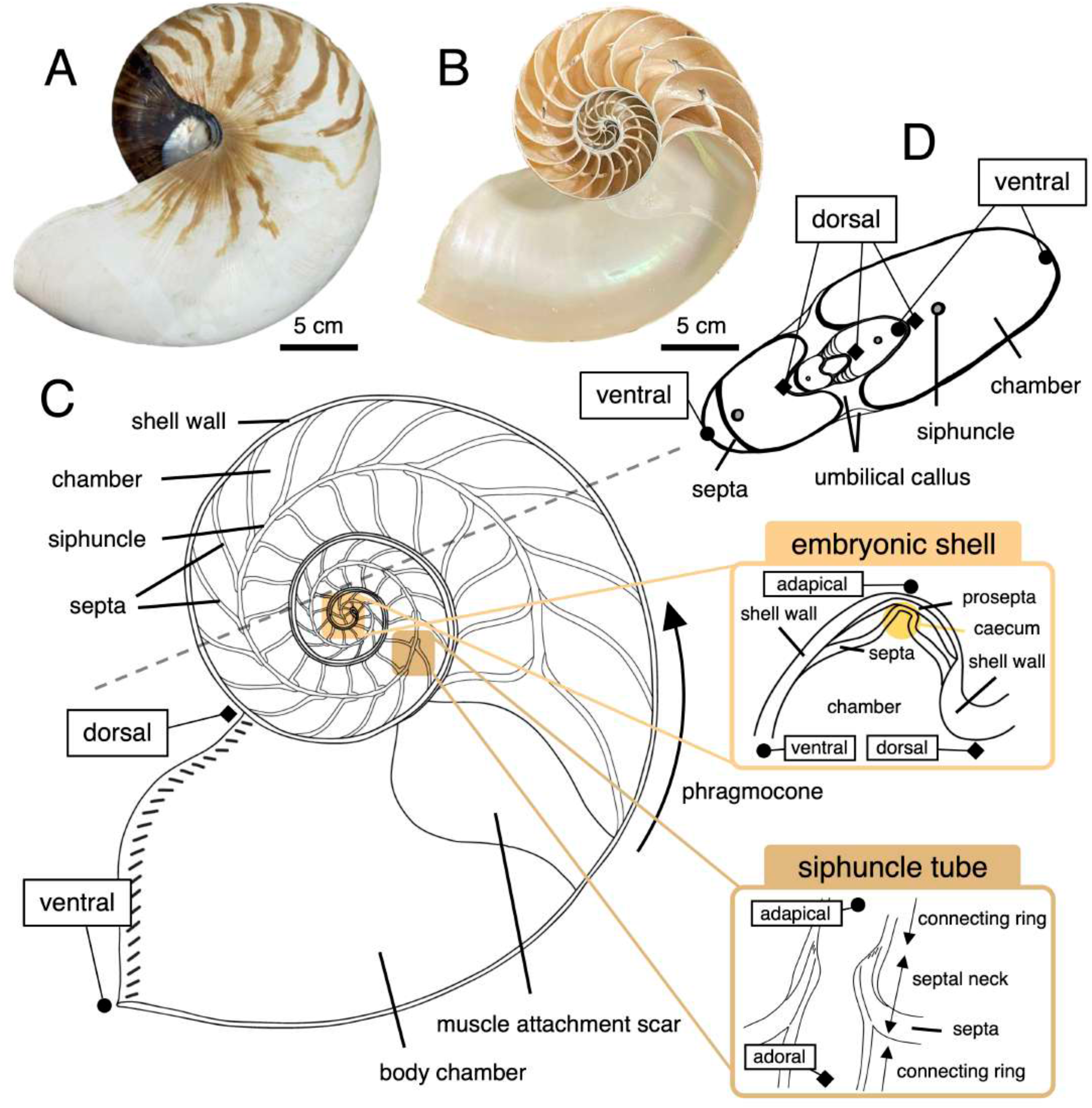
Structure and morphological terms for the nautilus shell. A–B) Photographs of the chambered nautilus *N. pompilius* and *N. cf. vitiensis*. A) Photograph of the whole shell in lateral view (specimen number: 20210301-036) and B) Horizontal section in lateral view. The scale bar represents 5 cm. C–D) Sketches of the nautilus shell components. C) Horizontal section in lateral view, and D) vertical section in lateral view. Enlarged views illustrate the embryonic shell region and the siphuncle–septum connection. Morphological and orientation terms are indicated in C. **Morphological technical terms**: body chamber, umbilical callus, chamber, connecting ring, muscle attachment scar, phragmocone, septa, septal neck, and siphuncle. **Morphological orientation terms**: adapical, adoral, dorsal, and ventral.

**Table 1.** Morphological glossary of the nautilus shell.

Like other cephalopods, nautilids undergo direct embryonic development without a metamorphic or larval transition between embryonic and adult stages (Arnold and Carlson 1986). Unlike other ectocochleate cephalopods, however, the nautilus lacks a protoconch, defined as the initial embryonic shell chamber separated from the subsequent phragmocone chambers (Drushchits 1977; Arnold and Carlson 1986). Therefore, the same shell that originates during embryogenesis continues to enlarge throughout life. Nautilus embryos undergo a developmental period of approximately eight to ten months before hatching, one of the longest among invertebrates (Saunders 1983; Tanabe and Uchiyama 1997; Shigeno et al. 2008). Shell secretion begins at around three months of embryonic development, producing the initial cup-shaped embryonic shell known as the cicatrix, marked by a black median pigment line that defines the coiling axis (Arnold and Carlson 1986; Landman and Cochran 2010; Sasaki et al. 2010). During the pre-chambered phase, the protoseptum is secreted as the first internal partition, followed by differentiation of the shell wall, septa, and siphuncle, forming the adult-type configuration before hatching (Landman et al. 1989; Tanabe and Uchiyama 1997). After hatching, the nautilus continues accretionary shell growth throughout its lifetime, eventually reaching an adult shell diameter of approximately 20 cm (Saunders 1983; Barord et al. 2023). Sexual maturity is typically reached around eight years of age, and individuals live for an additional five to ten years, with total lifespans often exceeding two decades (Saunders 1984b; Westermann et al. 2004; Dunstan et al. 2011b). In mature nautilus shells, growth decelerates and is accompanied by morphological modifications such as thickening of the apertural edge, development of a dark band along the inner aperture, and reinforcement of the final septum (Collins and Ward 2010).

The nautilus shell consists of three major components, the shell wall, septa, and siphuncle, together with several localized structures, such as the umbilical callus, hood attachment site, and muscle attachment scar, all formed by a complex interconnected microstructural system characterized by three-dimensional arrangements of aragonitic crystals and organic matrices (Grégoire 1962; Mutvei 1972; Landman et al. 1989; Mutvei and Doguzhaeva 1997; Tanabe and Uchiyama 1997; Arnold 2010). Like mollusk shells in general, the nautilus shell also has several localized structures associated with protection, reinforcement, attachment, and other functions (Thomas 1988). For example, the outer shell surface has two major localized features: the umbilical callus and the hood attachment site (Fig. 1; Table 1). The umbilical callus generally functions as a reinforcement structure covering the spiral center, and in *Nautilus* it is thickened and marked by a distinct black patch (Hirano and Obata 1979; Pietsch et al. 2021). Anterior to the umbilical callus, the hood attachment site is marked by dark deposits on the shell’s outer surface and provides the structural basis for sealing the aperture when the animal retracts (Owen 1832; Wells and Wells 1985). The animal’s soft body adheres to the shell through muscle cells, forming a scar known as the myostracum on the inner surface (Clark et al. 2020; Lee et al. 2011; Wu et al. 2017; Dong et al. 2022).

Despite this long history of research on *Nautilus* shell structure, most previous studies have focused on particular shell regions, developmental stages, or functional aspects. The shell has rarely been examined through an integrated morphomic framework that treats its major and localized components as parts of the same connected system. This leaves an important gap because the shell wall, septa, siphuncle, myostracum, embryonic shell region, apertural thickening, umbilical callus, and hood attachment site are not independent structures. Although they differ in position, timing of formation, morphology, and functional role, the components are formed within the same shell system and may share a constructional logic based on repeated use, modification, and connection of shell microstructures. Therefore, examining only one component or one shell region cannot fully clarify how different microstructures are distributed across the shell, how boundaries between components are formed, or how local modifications are added during ontogeny. In particular, the transitions between the shell wall, septa, and siphuncle are important because these interfaces connect structures involved in protection, chamber formation, buoyancy control, and mechanical reinforcement.

In this study, we conducted a scanning electron microscopy-based morphomic analysis (Son et al. 2023) of two vouchered museum shell specimens of *Nautilus*. We examined major shell components and localized structures, including the shell wall, septa, siphuncle, myostracum, embryonic shell region, apertural thickening, umbilical callus, and hood attachment site. By comparing these regions within and between specimens, we aimed to clarify the distribution of shell microstructural types, their transitions across component boundaries, and local variation associated with ontogeny and shell-region-specific modification. This approach allowed us to evaluate how microstructural heterogeneity among shell components and structural continuity across component boundaries are produced through repeated use, local modification, and connection of a limited set of shell microstructural types within the same shell system.

## Materials and Methods

### Specimen registration and collection information

In this study, we analyzed two legally obtained, vouchered museum shell specimens registered in the collection of the University Museum, The University of Tokyo (UMUT), specimen no. 20210301-036 of *Nautilus pompilius* (Linnaeus 1758), collected and imported from the Philippines in 1983, and specimen no. 20250701-001 of *Nautilus cf. vitiensis*, for which the collection date, import date, and locality are unknown (Fig. 1). The latter specimen was identified as *N. cf. vitiensis* based on shell coloration patterns, following Barord et al. (2023).

### Sample preparation and scanning electron microscopy (SEM) observation

Shell microstructures were observed using a scanning electron microscope (SEM; VE-8800, Keyence, Osaka, Japan). Shell samples were cut into smaller pieces with a precision cut-off machine (TS-45; Maruto Testing Machine Co., Tokyo, Japan). The observation surfaces were polished with sandpaper and treated with 1 M NaOH to remove residual organic matter. Polished sections were then etched with Mutvei’s solution for 15–30 min (Schöne et al. 2005). After ultrasonic cleaning for 10 min (USD-2R; AS ONE Corp., Osaka, Japan), samples were air-dried overnight. The surfaces were coated with osmium using an osmium coater (HPC-1SW; Vacuum Device Inc., Ibaraki, Japan), and SEM imaging was performed at an accelerating voltage of 10–15 kV.

### Measurement of the pigmented areas in the shell

Shell diameter at maturity and the pigmented shell area of the whole shell were measured following Barord et al. (2023). Photographs of the shells were edited using Adobe Photoshop v.24.7.5 (Adobe Systems, San Jose, California, USA). Subsequently, the pigmented areas of the whole shell were quantified using ImageJ v1.54g (Schneider et al. 2012).

### Measurement of the thickness of shell microstructures

To assess shell growth patterns at a microstructural level, the thickness of the shell wall layers was manually measured from SEM images of the *N. pompilius* specimen (UMUT 20210301-036). Two microstructural layers (an outer porcellaneous layer and an inner nacreous layer) of both the dorsal and ventral shell walls were measured. Landmarks were generally placed midway between septal attachment sites. The positions of the landmarks were determined by measuring the distances from the initiation point of the shell wall using ImageJ v1.54g (Schneider et al. 2012).

## Results

### 1. Five distinct microstructural types in the nautilus shell

We observed the presence of five microstructural types (spherulitic, prismatic, nacreous, semi-prismatic, and irregularly oriented prismatic structures; Fig. 2), identified based on crystal shape and orientation extending across layers of several tens of microns with only minor variations.

**Fig. 2.**
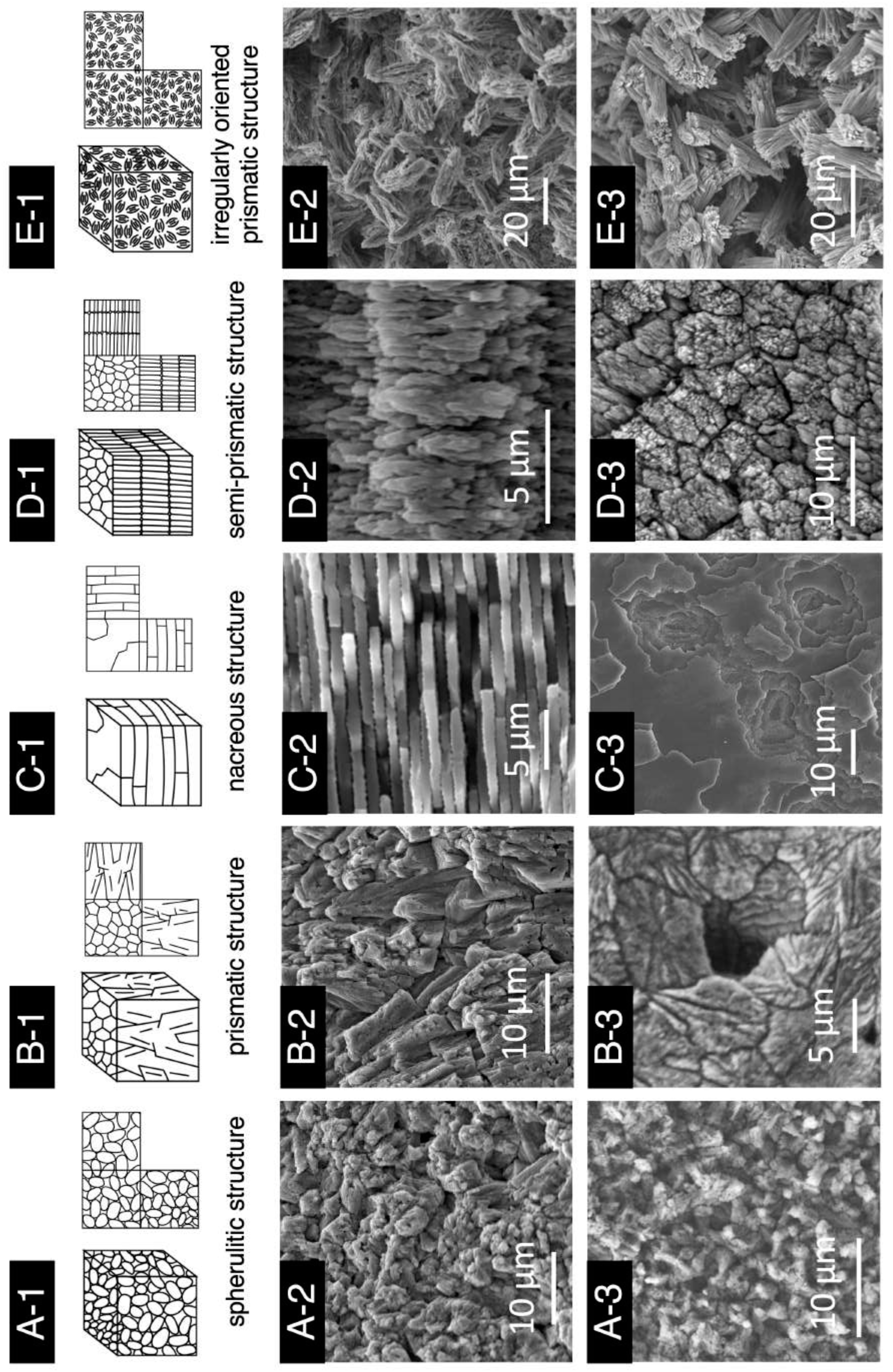
Five microstructural types in the nautilus shell. A–E) Microstructures of the nautilus shell: A) Spherulitic structure in the porcellaneous layer; B) Prismatic structure in the porcellaneous layer; C) Nacreous structure; D) Semi-prismatic structure; E) Irregularly oriented prismatic structure. 1) Three-dimensional drawing of each microstructure. 2–3) SEM images of the nautilus shell sections: 2) perpendicular to the shell surface and 3) parallel to the shell surface. B2 is a polished section, while the others are unpolished. A–D) Samples in A–D were etched for 15–30 min, whereas E was left unetched.

The spherulitic structure was characterized by granular crystals ca. 5 µm in diameter (Fig. 2A), while the prismatic structure was characterized by a prism-like structure that exhibited a unique triangular pyramid shape (Fig. 2B). Both microstructures were commonly observed side by side in the shell. In some regions, such as in those corresponding to early shell ontogeny, a radial growth of the prismatic structure from a center of nucleation (called spherulitic prismatic structure) was instead observed (e.g., Carter et al. 1989; Vinn and ten Hove 2011; Fig. 3C). The nacreous structure, the brick-wall configuration of mother-of-pearl, exhibited a layered arrangement composed of stacks of tablets, which were organized in lamellae separated by interlamellar conchiolin membranes at intervals of ca. 1.5 µm (Fig. 2C). These tabular crystals of the nacreous layer were formed by stacking on top of preceding lamellae, grew laterally, and extended until they contacted adjacent stacks, creating a tightly aligned side-by-side arrangement.

**Fig. 3.**
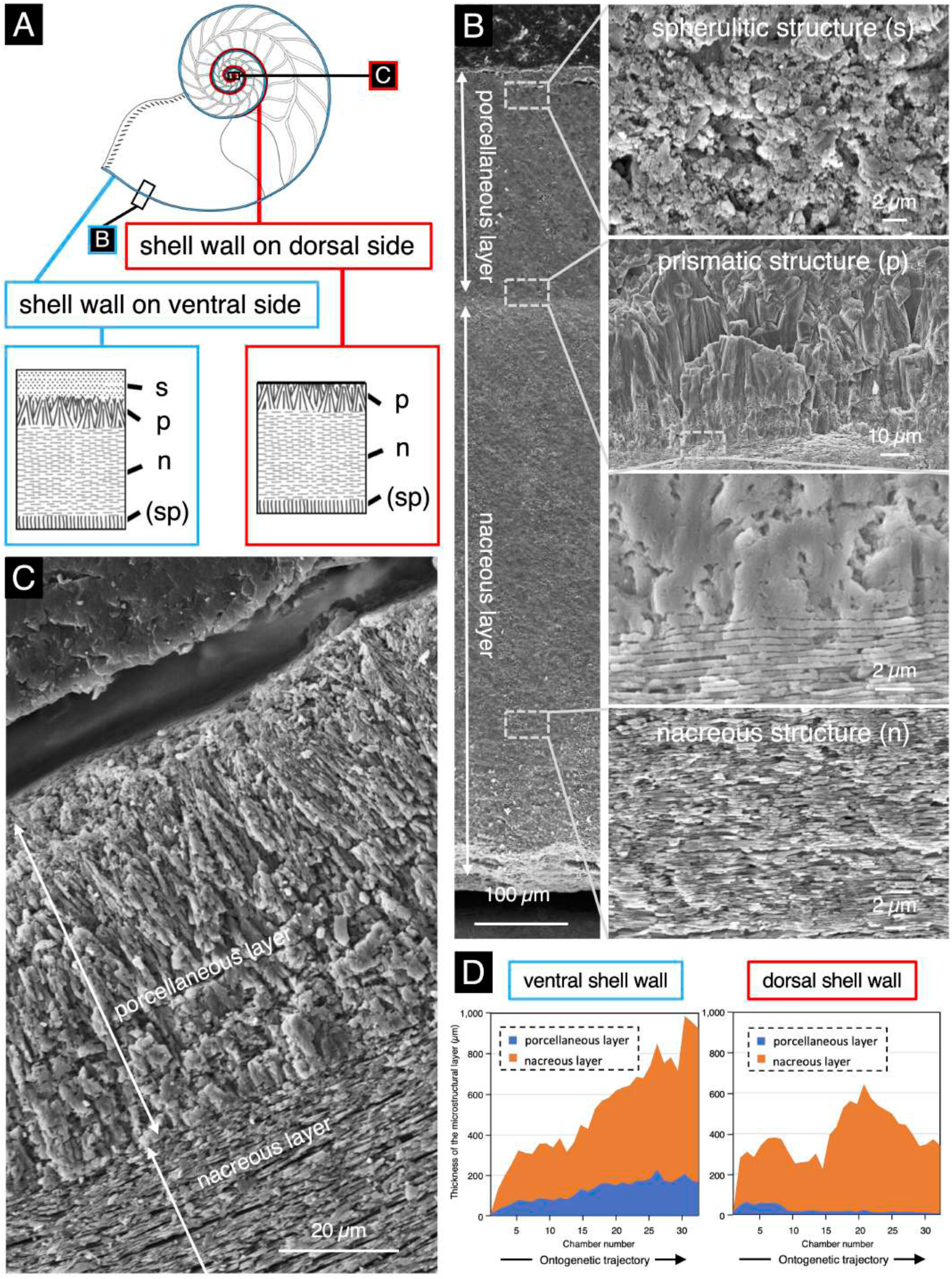
Microstructure of the shell wall. Microstructure of the shell wall of the nautilus shell. A) Schematic diagrams of the microstructure of the shell wall. The colored regions indicate the ventral (blue) and dorsal (red) sides of the shell wall. B, C) SEM images of the microstructure of the ventral (B) and dorsal (C) shell walls. D) Ontogenetic changes in the thicknesses of the porcellaneous and nacreous layers. **Abbreviations**: spherulitic structure (s), prismatic structure (p), nacreous structure (n), semi-prismatic structure (sp)

The fourth type of microstructure, the semi-prismatic structure, exhibited rounded, columnar crystal units, intermediate in form between needles and prisms, arranged perpendicular to the shell surface (Fig. 2D). The term semi-prismatic structure was originally defined by Mutvei (1964), and his later paper replaced it with “prismatic structure” because of limitations in detailed characterizations (Mutvei 1972). However, to clearly distinguish the morphologically different prismatic structures in our observation, we resurrected the term semi-prismatic structure. This term was introduced by Mutvei (1964) but later subsumed under “prismatic structure” (Mutvei 1972), probably because that later usage was based on observations from a limited shell region and therefore did not capture the full diversity of prismatic structures in the nautilus shell (Fuchigami and Sasaki 2005; Vinn 2013; Sato and Sasaki 2015; Checa 2018).

The last type of microstructure observed in the nautilus shell was the irregularly oriented prismatic structure. The structure was characterized by sheaf-like spherulites (ca. 20 µm in height and 5 µm in width) with isotropic and irregular orientation (Fig. 2E). In previous observations of the Nautilus shells, the irregularly oriented prismatic structure was referred to as “spherulitic prismatic structure” (e.g., Mutvei 1972; Tanabe 1982), but this term was ambiguous and overlaps with another definition described above (e.g., Carter et al. 1989; Dauphin et al. 2020). Therefore, in this study, we adopted the term “irregularly oriented prismatic structure” instead (Vinn and ten Hove 2011).

Further observations showed that the arrangement and distribution of these microstructures vary across different shell components. The spherulitic and prismatic structures were generally found in the outer regions of the shell wall, whereas the nacreous structure dominated its inner parts (Mutvei 1972; Tanabe 1982; Fig. 2F). The semi-prismatic structure was locally developed in transition zones between the nacreous layers of the shell wall and the septum (Mutvei 1972; see Sections 2.1 and 2.4; Fig. 3A), and between the nacreous layer of the septum and organic layer of siphuncle (see Section 3.1; Fig. 7D–F, H). The irregularly oriented prismatic structure was observed in limited areas, often occurring near growth boundaries or along structural connections (Tanabe 1982; Fig. 3B). The occurrence and relative proportions of these five microstructures differed slightly among specimens and shell regions, indicating localized variation in crystal organization during shell growth (Fig. 3C).

### 2. Distinct combinations of the five microstructural types among major morphological components of the nautilus shell

We conducted stereomicroscopic and scanning electron microscopy (SEM) observations on mature nautilus shell specimens (Figs. 3–6) to examine how the five microstructural types (spherulitic, prismatic, nacreous, semi-prismatic, and irregularly oriented prismatic) and associated organic matrices were distributed in the main morphological regions of the shell. The results showed that each shell component exhibited a distinct structural combination of these microstructural types (Fig. 3).

**Fig. 4.**
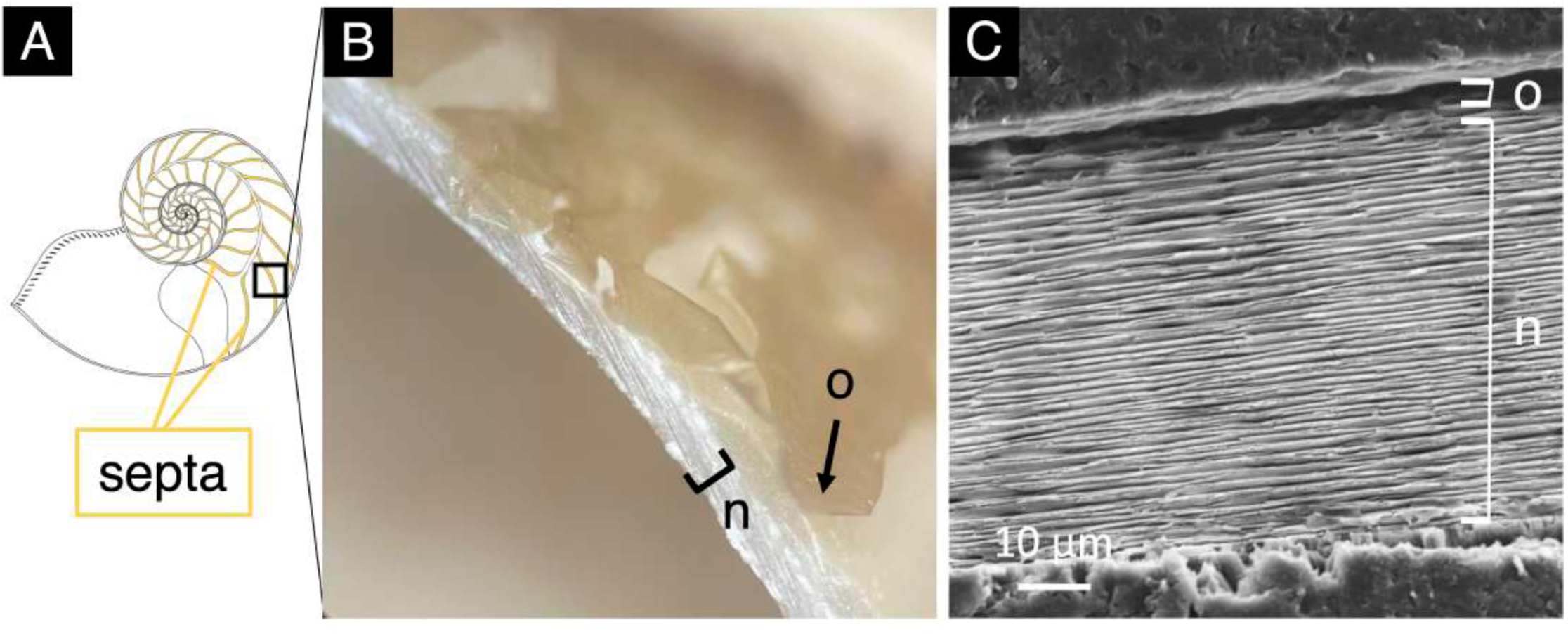
Microstructure of the septum. A, B) Schematic diagram (A) and photograph (B) of the septum. C) SEM image of the microstructure of the septum. **Abbreviations**: organic layer (o), nacreous layer (n)

**Fig. 5.**
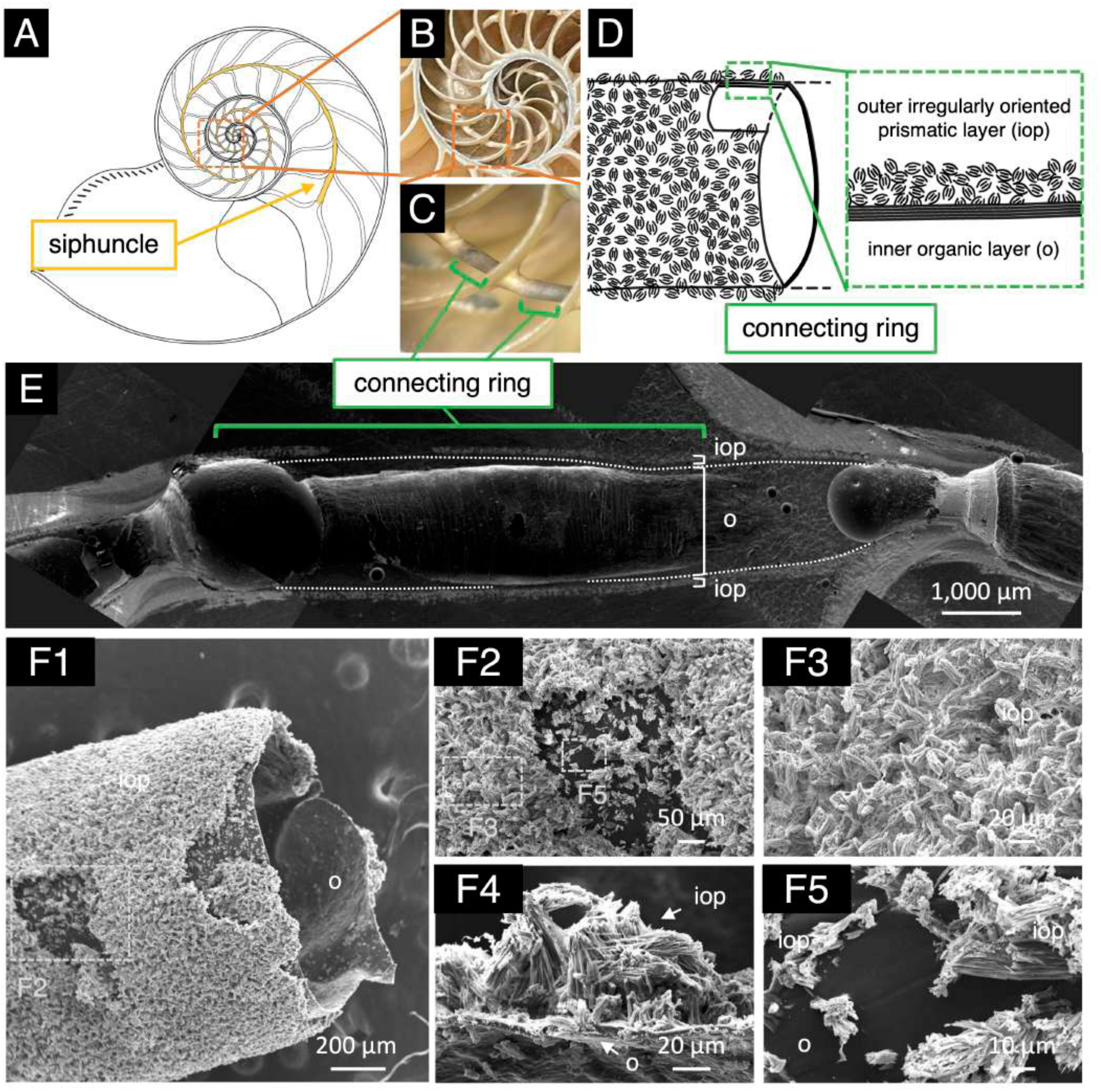
Microstructure of the connecting ring of the siphuncle. A) Schematic diagram of the connecting ring of the siphuncle. B, C) Photographs of the connecting ring under dry (B) and wet (C) conditions. D) Schematic diagram of the microstructure of the connecting ring. E, F) SEM images of the microstructure of the connecting ring: (E) section embedded in epoxy resin and (F) unetched section. **Abbreviations**: irregularly oriented prismatic structure (iop), organic structure (o).

**Fig. 6.**
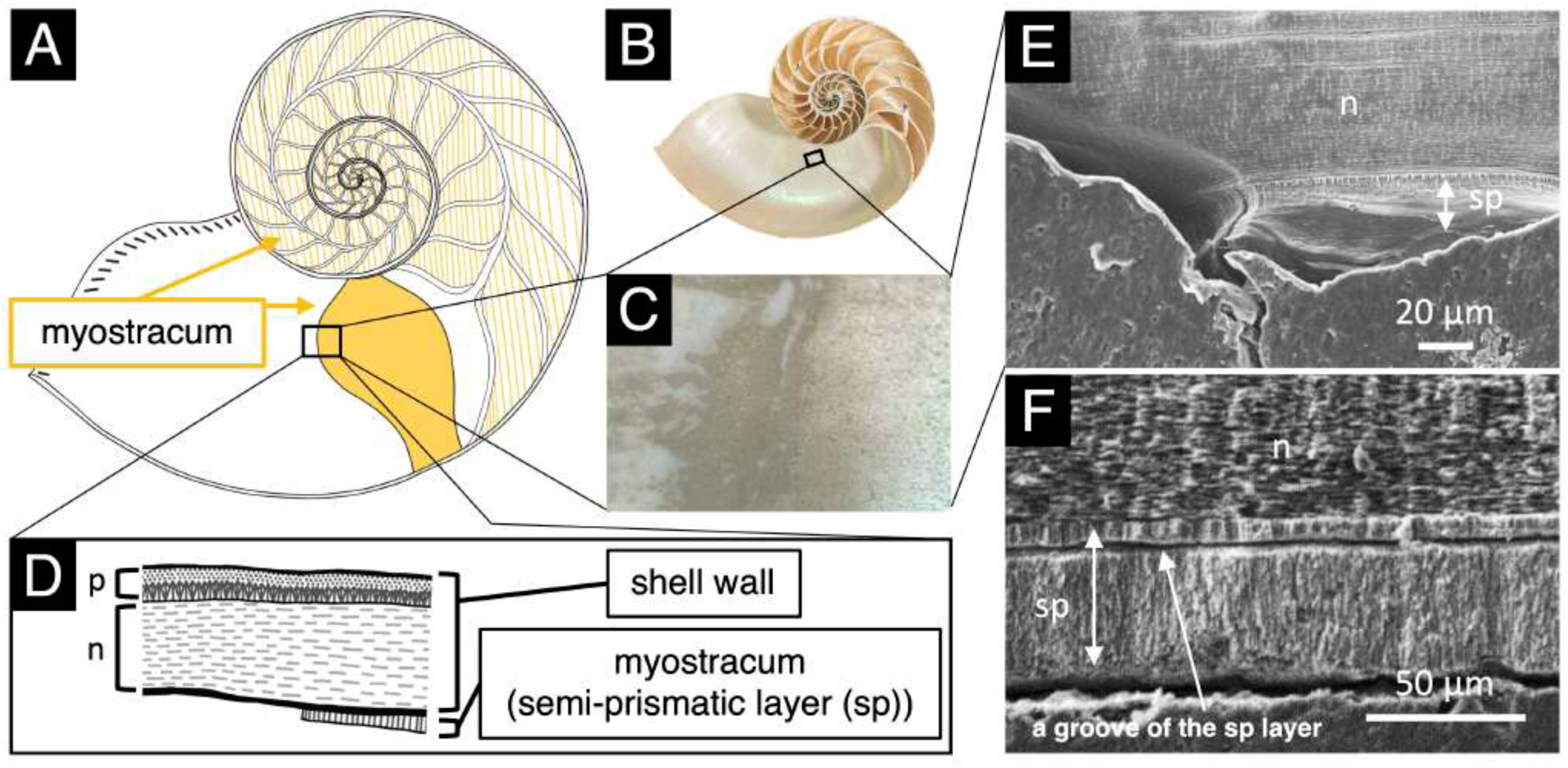
Microstructure of the myostracum at the muscle attachment site. A) Schematic diagram of the distribution of the myostracum. B, C) Photographs of the muscle scar. D) Schematic diagram of the microstructure of the myostracum layer. E, F) SEM images of the myostracum layer. (E) unetched and (F) heavily etched sections. Abbreviations: porcellaneous structure (p), nacreous structure (n), semi-prismatic structure (sp)

#### 2.1. Shell Wall

The shell wall forms the external component of the shell and exhibits orange-colored stripes on a generally white area ventrally (Figs. 1, 3A). Its exterior surface is rough, while the interior surface is smooth and glossy. Based on the SEM observations, the shell wall consisted of two layers: an outer porcellaneous layer (spherulitic–prismatic layer) and an inner nacreous layer (Fig. 3B). The porcellaneous layer, which corresponds to the rough exterior surface, consisted of two sub-layers: an outer spherulitic layer and an inner prismatic layer (Figs. 2A, B, 3B). In contrast, the nacreous layer, which corresponds to the glossy interior surface, was characterized by a brick-and-mortar structure formed by the arrangement of nacreous tablets (Figs. 2C, 3B, C). In addition to these two layers, an innermost semi-prismatic layer was observed in the region of the myostracum representing the muscle attachment (see Section 2.4; Fig. 6).

Three microstructures of the shell wall (spherulitic, prismatic, and nacreous) could be distinguished by their crystal morphologies. However, the boundaries between them were not clear. The interfaces formed transitional zones characterized by a gradual transition from the spherulitic to the prismatic, and from the prismatic to the nacreous arrangements (Figs. 3B, C). The spherulitic–prismatic transition occurred progressively, in which the prismatic structure was initiated at the center of nucleation of the spherulitic structure (Fig. 3B). At the boundary between the prismatic and nacreous structures (Fig. 3B, C), interlamellar conchiolin membranes characteristic of the nacreous structure began to emerge fragmentarily at the edge of the prismatic structure, indicating an abrupt shift in microstructural organization.

The position of the transition between the prismatic and nacreous structures varied spatiotemporally, reflecting ontogenetic and dorsoventral differences in the thickness of the outer porcellaneous layer and the inner nacreous layer (Fig. 3B–D). In the ventral shell wall, the thickness of both the outer porcellaneous and the inner nacreous layers increased linearly throughout ontogeny, from embryonic development through maturity (Fig. 3D). In contrast, the two layers in the dorsal shell wall did not exhibit a linear pattern of thickening. That is the thickness of the outer porcellaneous layer was largely constant, with a marked difference between the embryonic stage (mean = 53.0 µm, SD = 6.44, for chambers 2–9, i.e., the first whorls) and the post-hatching stage (mean = 14.9 µm, SD = 3.39, for chambers 9–31). The outer porcellaneous layer gradually thinned adorally after the 31st chamber interval and eventually disappeared. As for the nacreous layer, the thickness was nearly constant until the 15th chamber (mean = 231 µm, SD = 34.1, for chambers 2–15), then increased up to the 22nd chamber (ca. 599 µm), and thereafter gradually decreased toward the final chambers.

Consistent with this ontogenetic growth pattern, the thickness of the shell wall differed spatially along the dorsoventral axis. For instance, at the 31st septal attachment region, the ventral shell wall reached 778.7 µm in thickness (208.7 µm for the outer porcellaneous layer and 570.0 µm for the inner nacreous layer; Fig. 3B, D). In contrast, the dorsal shell wall at the same septal attachment region was less than half as thick, measuring 336.7 µm (10.0 µm for the outer porcellaneous layer and 326.7 µm for the inner nacreous layer; Fig. 3C, D).

#### 2.2. Septum

The septa internally partition the shell into chambers (Figs. 1, 4A). They form concavely curved surfaces facing the adoral side, with more than 30 present in mature individuals (*N*. *pompilius* 20210301-036: 31 septa; *N. cf. vitiensis* 20250701-001: 33 septa). A fractured section observed under a stereomicroscope showed a thin organic layer on the adapical side that was peeled away from the nacreous layer (Fig. 4B). SEM images showed that the septum primarily consisted of a nacreous layer (Figs. 2C, 4C). Each septum was connected to the shell wall along the outer margin and to the siphuncle around the central opening, with an additional semi-prismatic layer present at both connections (see Sections 3.1 and 3.2).

#### 2.3. Connecting ring of the siphuncle

The siphuncle, a tube that penetrates the septa in the center, functions as a permeable channel for the body fluid within newly formed chambers, regulating buoyancy and contributing to mechanical strength (Figs. 1, 5A). The connecting ring corresponds to the tubular portion between septal connections. Stereomicroscopic observations revealed that the structure consisted of a calcified layer overlying a black organic layer, and it was thin and fragile (Fig. 5B–D). Because of its fragile properties, the connecting ring was inadvertently destroyed during treatment, including resin embedding and etching (Fig. 5E). Untreated sections showed the microstructure more clearly, with less damage (Fig. 5F). The microstructure of the connecting ring consisted of two layers: an outer irregularly oriented prismatic layer and an inner organic layer (Fig. 5D–F). The outer prismatic layer was thin (ca. 80 µm) and porous. Its crystals formed complexes with organic matter and grew upon the inner organic layer (Fig. 5E, F). This prismatic layer extended into the septum at the septal neck and was connected to a semi-prismatic (pillar-like) layer (see Section 3.1). The inner organic layer was wrinkled along the longitudinal axis of the tube and lacked porosity (Fig. 5F). This inner layer extended into the septum at the septal neck and became segmented into other organic portions within the septal neck region (see Section 3.1).

#### 2.4. Myostracum

The muscle scar, or myostracum, is the site where muscle cells attach to the shell, and these attachment areas are visibly marked on the inner surface of the shell (Figs. 1, 6A–C). The microstructure of the muscle attachment site was characterized by a semi-prismatic layer (referred to as a myostracum layer; Dong et al. 2022) (Fig. 6D–F). The crystal morphology of this layer was clearly observed near its connection with the shell wall, but it became indistinct toward the interior surface of the muscle attachment site (Fig. 6E). In highly etched sections, the semi-prismatic layer of the myostracum was separated into two distinct regions (Fig. 6F). An intervening groove appeared between these two distinct regions, indicating the presence of organic-rich material, as the etching process preferentially removed organic components (Fig. 6E, F). This groove within the semi-prismatic layer served as one of the sites from which septal nacreous layers originated (see Section 3.2).

### 3. Shell component connections at transitions between distinct microstructures

We conducted scanning electron microscopy (SEM) observations of connections between shell components (Figs. 7–8) and identified two types of junctions: the siphuncle–septum junction (septal neck) and the shell wall–septum junction, each showing structural transitions between the microstructures of adjoining components.

**Fig. 7.**
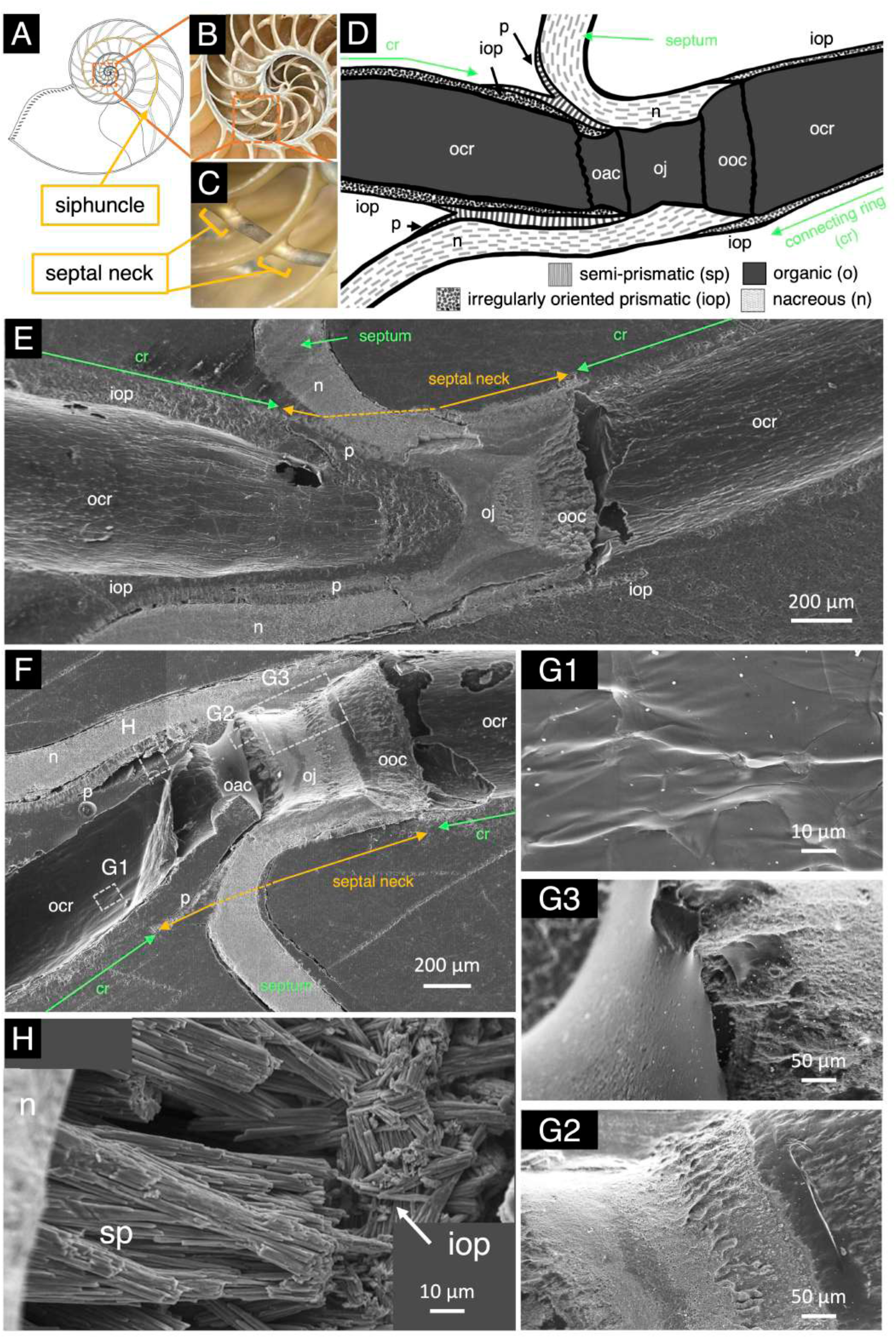
Microstructure of the septal neck of the siphuncle. A) Schematic diagram of the septal neck. B, C) Photographs of the septal neck. D) Schematic diagram of the microstructure of the septal neck. E–H) SEM images of the microstructure of the septal neck. E, F) Overview images from a parasagittal section (E) and a vertical section (F). G, H) Enlarged views of panel F, showing the organic segments (G1–G3) and the prismatic structure surrounding the septal neck (H). **Abbreviations**: nacreous structure (n), irregularly oriented prismatic structure (iop), semi-prismatic structure (sp), organic connecting ring segment (ocr), organic adapical cap segment (oac), organic adoral cap segment (ooc), organic joint segment (oj)

**Fig. 8.**
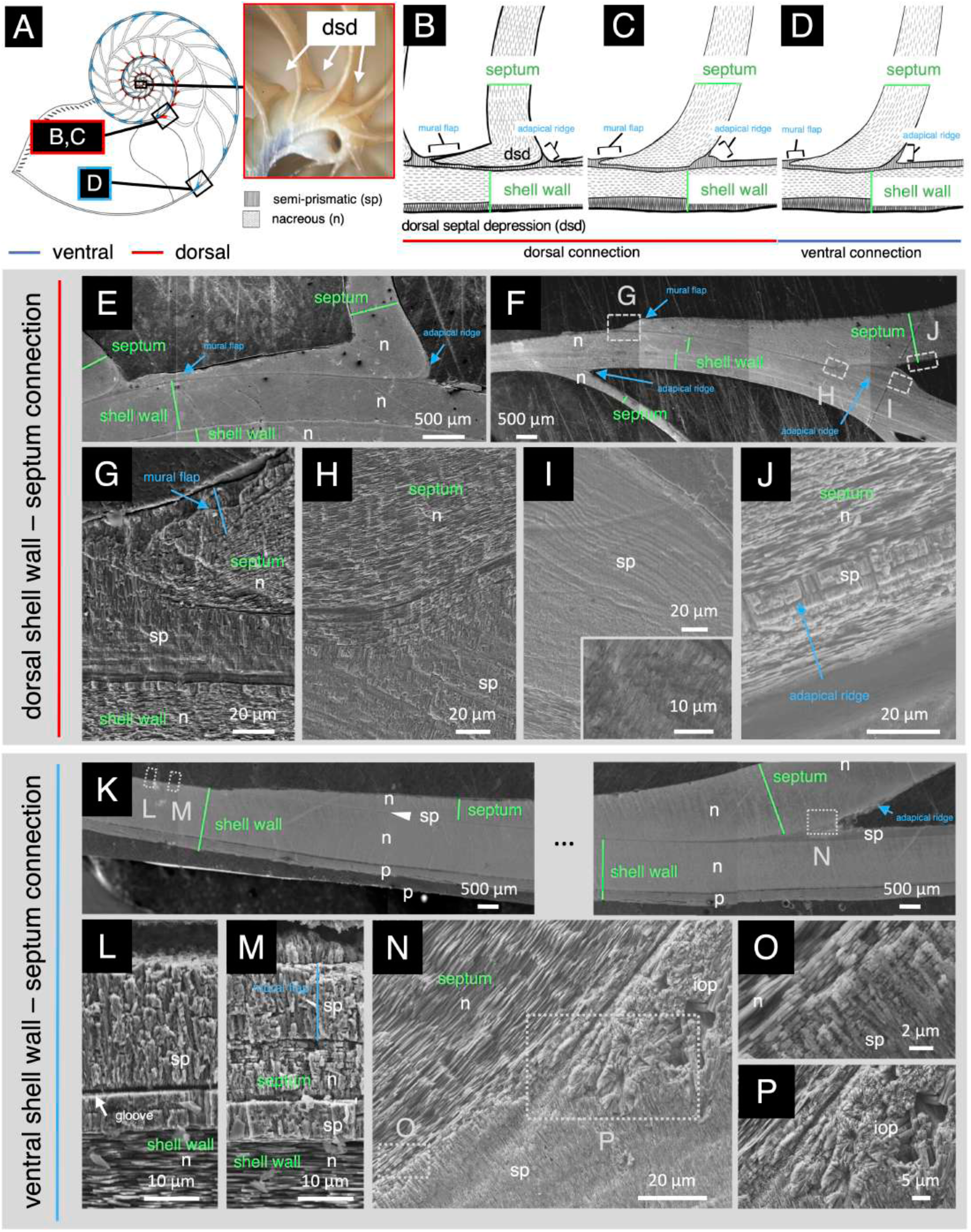
Microstructure of the shell wall–septum connection. A–D) Schematic diagrams of the microstructure of the shell wall–septum connection across ontogeny: A) Position of the shell wall–septum connection, B) the dorsal connection with a dorsal septal depression (dsd), C) the dorsal connection without a dsd, D) the ventral connection. E–N) SEM images of the microstructure at the shell wall–septum connection. E–J) The dorsal connection; G–J are enlarged views of F. K–N) The ventral connection; L–N are enlarged views of K. O and P are enlarged views of N. **Abbreviations**: prismatic structure (p), nacreous structure (n), semi-prismatic structure (sp), irregularly oriented prismatic structure (iop).

#### 3.1. Siphuncle–septum connection (Septal neck)

The septal neck is a part of the siphuncle that connects the siphuncle to the septum (Fig. 7A–C). SEM images showed that the septal neck consisted of an organic layer and three calcified layers: a nacreous layer, a semi-prismatic layer, and an irregularly oriented prismatic layer (Fig. 7D–G). The organic layer and the irregularly oriented prismatic layer were derived from the connecting ring of the siphuncle, while the nacreous layer was derived from the septum. The semi-prismatic layer was specifically observed in the septal neck.

The organic layer in the septal neck was clearly visible and could be subdivided into four organic segments based on shape and texture: a connecting ring (ocr), an adapical cap (oac), an adoral cap (ooc), and a joint (oj) (Fig. 7D–F). The connecting ring segment extended from the root of the septal neck toward the tip of the subsequent septal neck, while the other organic segments (the adapical cap, the adoral cap, and the joint) were restricted to the septal neck. The adapical and adoral cap segments were located at the adapical and adoral ends of the connecting ring segment in the septal neck and formed connective sleeves with trapezoidal outlines (Fig. 7F, G). The adapical cap segment exhibited a smooth surface, while the adoral cap segment exhibited a rugged surface (Fig. 7G2, G3). The adapical cap segment was detached from the septum and appeared deformed, likely as a result of damage during the epoxy embedding. The joint segment connected adjacent connecting rings through the adapical and adoral cap segments (Fig. 7F; also observed in the *Spirula* shell; Mutvei 2017).

The nacreous layer of the septum connected to both the joint and the adoral cap segments (Fig. 7D–F). The abundance of the interlamellar conchiolin membranes of the nacreous structure gradually increased toward the siphuncular connection and eventually predominated at the joint. Therefore, the boundaries between the nacreous layer and the organic layer in the septal neck became indistinguishable, as the two organic structures are fully integrated.

The semi-prismatic layer grew from the interior surface of the septal nacreous layer, around the septal neck (Fig. 7D–F, H). This layer was connected to the root of the joint segment. At the site where this layer detached from the siphuncle, it was observed as a pillar-like structure (referred to as the pillar layer or pillar zone in *Spirula* shell; Bandel 1989; Bandel and Boletzky 1979; Mutvei 2017) (Fig. 7H). This pillar layer is continuous with the outer irregularly oriented prismatic layer of the connecting ring, showing a gradual increase in the disorder of crystal arrangement across the transition (Fig. 7H). The irregularly oriented prismatic structure of the connecting ring further extended into the nacreous layer of the subsequent septal neck (Figs. 5D, 7H).

#### 3.2. Shell wall–septum connection

The connection between the shell wall and the septum forms a key architectural interface within the nautilus shell (Fig. 8). The morphology of this interface was not uniform but exhibited marked dorsoventral differentiation during shell ontogeny (Fig. 8B–E). At the dorsal connection, the septum formed a locally developed concavity toward the adapical side, referred to as the dorsal septal depression (dsd; Fig. 8C; Mutvei and Doguzhaeva 1997). The depth of the dsd progressively increased beginning around the 7th septum, reached its maximum in the septa of the 2nd whorl, and then became less distinct again in the septa of the 3rd whorl (Mutvei and Doguzhaeva 1997). Corresponding to this ontogenetic pattern, SEM images showed that the vertical sections of the dorsal septum exhibited a dogleg-like shape in early stages (Fig. 8B, E), but became more shallowly curved as the dsd diminished (Fig. 8C, F). Simultaneously, septal spacing also increased (Fig. 8A). In contrast, the ventral connection consistently exhibited a shallow curvature and relatively uniform septal spacing (Fig. 8D, K). These dorsoventral differences were also observed in the shell wall during early development (see Section 4.1).

Both the shell wall and septum primarily consisted of nacreous layers (Sections 2.1 and 2.2), but these nacreous layers were completely independent, being separated by the myostracum, a semi-prismatic layer (Fig. 8; see Section 2.4). The nacreous layer of the septum originated from a region within the semi-prismatic layer (Fig. 8L, M). Crystals in the semi-prismatic layer were generally elongated perpendicular to the shell surface (Fig. 8B–D, G–J, L–N). The semi-prismatic structure surrounded the septal nacreous layer and could be subdivided into three regions: a thin region sandwiched between two nacreous layers (∼10 µm in thickness), a mural flap, and an adapical ridge flap (corresponding to the region referred to as cameral deposits in the paleontological literature; Fischer and Teichert 1969; Pohle et al. 2025) (Fig. 8B–D; Lemanis et al. 2020; Checa et al. 2022). The mural flap overlaid the internal face of the septum at the adoral edge, while the adapical ridge flap overlaid the septum at the adapical edge (Fig. 8B–D). Both flaps represented localized thickenings of the semi-prismatic layer at the shell wall–septum connection, where the semi-prismatic structure formed a distinct striped pattern (Fig. 8G–I, L–N). The shell wall–septum connection, including these overlying flaps, showed continuous transitions between semi-prismatic and nacreous structures on both sides of the interface (Fig. 8G, H, M, O). Moreover, in the adapical ridge region not directly connected to the nacreous layer, the semi-prismatic structure transitioned into an irregularly oriented prismatic structure, resembling that of the septal neck (Fig. 8P).

### 4. Microstructural observations of shell ontogeny

We conducted scanning electron microscopy (SEM) observations of shell regions corresponding to early and late ontogenetic stages in two adult shells (Figs. 9–10). Shell regions formed early in ontogeny preserve the initial condition of the shell wall, septum, and siphuncle, representing the earliest stage of shell construction recorded in the shells (Fig. 9). In these regions, the caecum, as the initial part of the siphuncle, varied between specimens (Fig. 9). In contrast, shell regions formed later in ontogeny exhibited mature microstructural organization (Fig. 10).

**Fig. 9.**
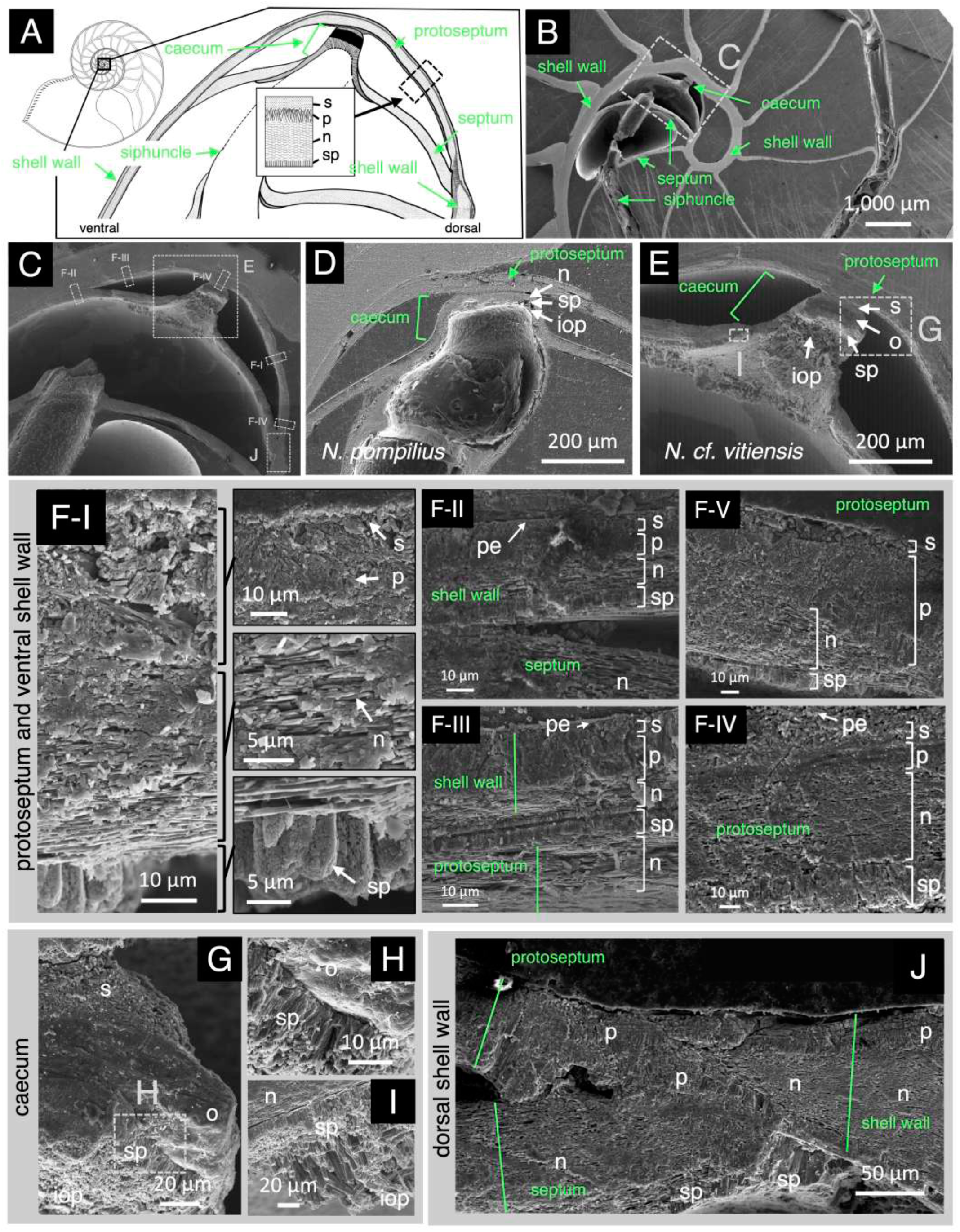
Microstructural organization of the embryonic shell region. A) Schematic diagram of the microstructure of the *Nautilus cf. vitiensis* embryonic shell. B–J) SEM images of the microstructure of the nautilus embryonic shell. B, C) SEM overviews of the embryonic shell microstructure; C is an enlarged view of B. D, E) Enlarged views of the protoseptal microstructure of *N. pompilius* (D) and *N. cf. vitiensis* (E). F) Microstructure of the protoseptum and ventral shell wall; F is an enlarged view of C. G–I) Microstructure of the caecum of *N. cf. vitiensis*; G and I are enlarged views of E. J) Microstructure of the dorsal shell wall; J is an enlarged view of C. **Abbreviations**: spherulitic structure (s), prismatic structure (p), nacreous structure (n), semi-prismatic structure (sp), irregularly oriented prismatic structure (iop).

**Fig. 10.**
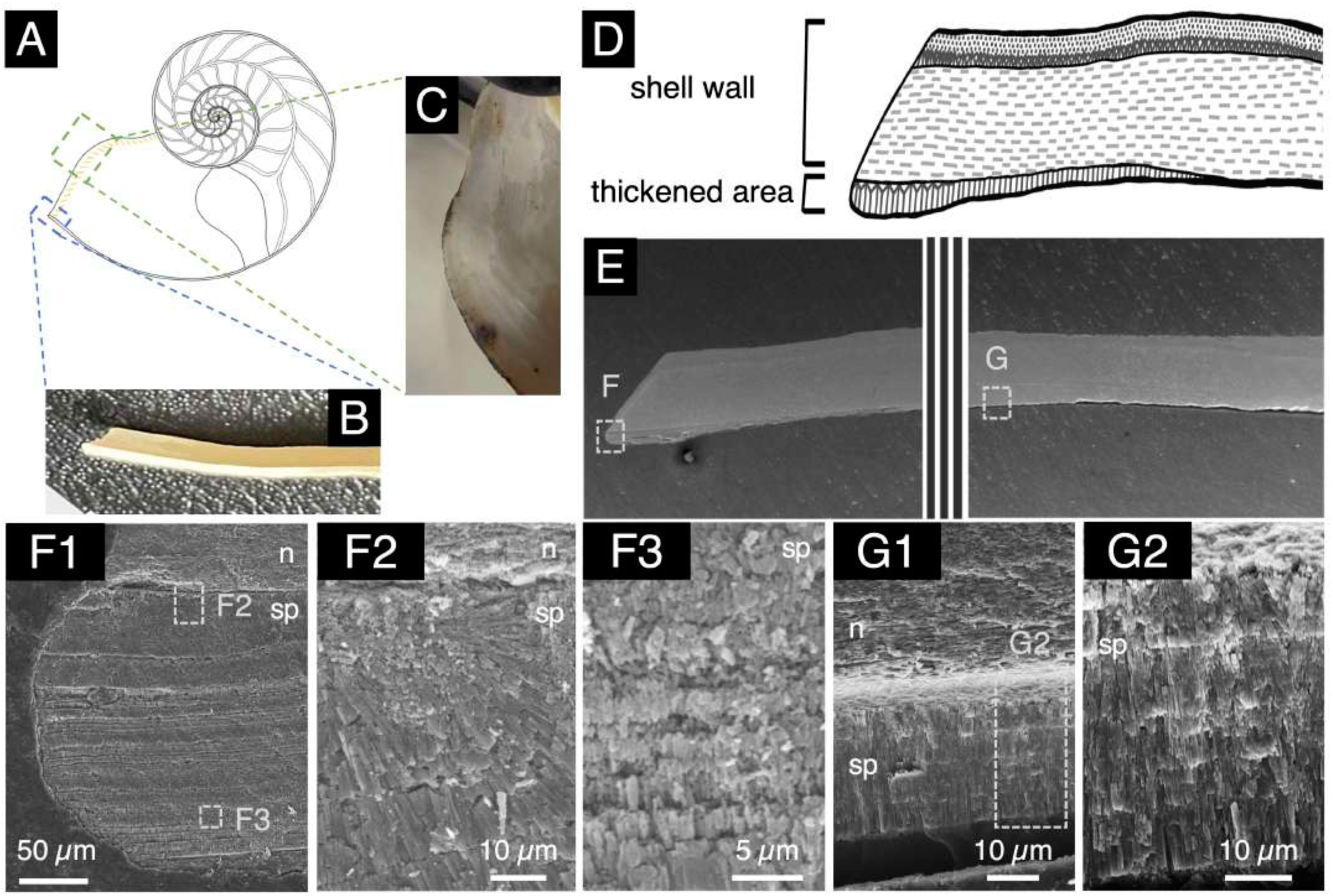
Microstructure of shell-aperture thickening. A) Schematic diagram of the thickened area of the shell aperture. B, C) Photographs of the thickened area of the shell aperture: B) vertical view; C) surface view. D) Schematic diagram of the microstructure of the thickened area of the shell aperture. E–G) SEM images of the thickened area of the shell aperture. F and G are enlarged views of E.

#### 4.1. Early shell formation during embryonic development

In embryonic shell development, the nautilus first formed the cicatrix, followed by both the shell wall and the protoseptum (Tanabe and Uchiyama 1997; Saunders and Landman 2009). The cicatrix was not observable in this study, as this region was too small to detect in vertical sections. Instead, the protoseptum represented the outermost calcified layer of the embryonic shell (Figs. 1, 9A, F). This layer exhibited a rudimentary state characterized by the intermingling of spherulitic and prismatic structures, which gradually differentiated into the porcellaneous layer of the shell wall, with no distinct boundary between the two. A nacreous layer subsequently developed beneath this rudimentary layer as part of the protoseptum. This nacreous layer was not continuous with that of the shell wall, gradually thinning and terminating before reaching the shell-wall nacreous layer (Fig. 9A, F). The protoseptum constituted the apical surface of the first chamber but was not a true septum, as it was not penetrated by the siphuncle. Furthermore, an inner semi-prismatic layer was present beneath the nacreous layer, and the structure resembled the pillar-like prismatic structure observed in the septal neck, rather than the semi-prismatic structure of the myostracum. These observations indicate that, following the formation of the primordial shell wall represented by the outer rudimentary layer, the protoseptum was established as an initial septum before the development of the siphuncle tube.

Following the formation of the rudimentary outer layer of the protoseptum, the nacreous layer of the shell wall began to develop prior to the completion of the protoseptum. The development pattern of the shell-wall nacreous layer differed between the ventral and dorsal regions. In the ventral region, the nacreous layer of the shell wall originated by splitting from the outer rudimentary layer, which subsequently remained positioned between the outer layer of the shell wall and that of the protoseptum (Figs. 9F, J, S2). The nacreous layer gradually increased in thickness following its formation, whereas the intervening rudimentary layer was continuous with the myostracum layer. In contrast, the dorsal shell wall formed abruptly at the point where the nacreous layer of the protoseptum completely disappeared, and its thickness increased rapidly (Figs. 9F, J, S2). These observations revealed contrasting modes of nacreous-layer formation between the ventral and dorsal shell walls.

The caecum was a sac-like initial part of the siphuncle (Fig. 9A). It developed in connection with the semi-prismatic structure of the protoseptum (Fig. 9D, E, G–I). The caecum was composed of spherulitic, nacreous, irregularly oriented prismatic, and organic structures (Fig. 9D, E, G–I). Notably, the caecum exhibited different microstructural organizations in the two specimens examined, *N. pompilius* (ID: 20210301-036) and *N. cf. vitiensis* (ID: 20250701-001), the latter being assigned based on morphological characteristics (Fig. S1; Barord et al. 2023) (Fig. 9C–E). In *N. pompilius* the caecum was formed by a curvature of the septum, which consisted of the outermost organic layer and an outer nacreous layer (Fig. 9D). On the adoral side of the curved septum, rudimentary pillar-like prismatic structures and irregularly oriented prismatic structures were observed together, forming the attachment site for the organic layer of the siphuncle (Fig. 9D). In contrast, the caecum of *N. cf. vitiensis* exhibited a distinct morphology arising from a localized curvature of the septal nacreous layer (Fig. 9E). It was composed of an outer spherulitic layer, a thin middle organic layer, and an inner semi-prismatic layer (Fig. 9E, G–I). Toward the tip of the caecum, the septal nacreous layer gradually decreased in thickness and became indistinguishable from the outer spherulitic layer. The middle organic layer was distinct from the siphuncular organic layer. Internally, the semi-prismatic layer gradually transitioned into an irregularly oriented prismatic structure, which served as the attachment site for the siphuncular organic layer, resembling that of the septal neck. The structures contributing to caecum formation differed markedly between the two specimens, involving different combinations of microstructures and organic layers.

#### 4.2. Apertural thickening during maturation

The shell aperture was thickened in mature specimens (Fig. 10). The thickened region corresponded to an additional layer coating the inner surface of the shell wall, which exhibited black patches and a slightly rough texture (Fig. 10C). This layer consisted of a semi-prismatic structure resembling that of the myostracum (Fig. 10F, G). In some regions, the semi-prismatic structure showed radial growth on the nacreous layer of the shell wall, with semi-prismatic crystals growing successively on the underlying crystals (Fig. 10F, G). The thickness of this semi-prismatic layer reached a maximum of approximately 220 µm near the distal shell aperture and gradually decreased proximally.

### 5. Appendage parts of the exterior surface of the nautilus shell

We conducted scanning electron microscopy (SEM) observations to examine the umbilical callus and the hood attachment site, both of which constitute appendage parts of the exterior surface of the nautilus shell (Figs. 11–12). These structures were not formed by unique shell microstructures but were constructed using microstructure types and organic layers observed in other shell components.

**Fig. 11.**
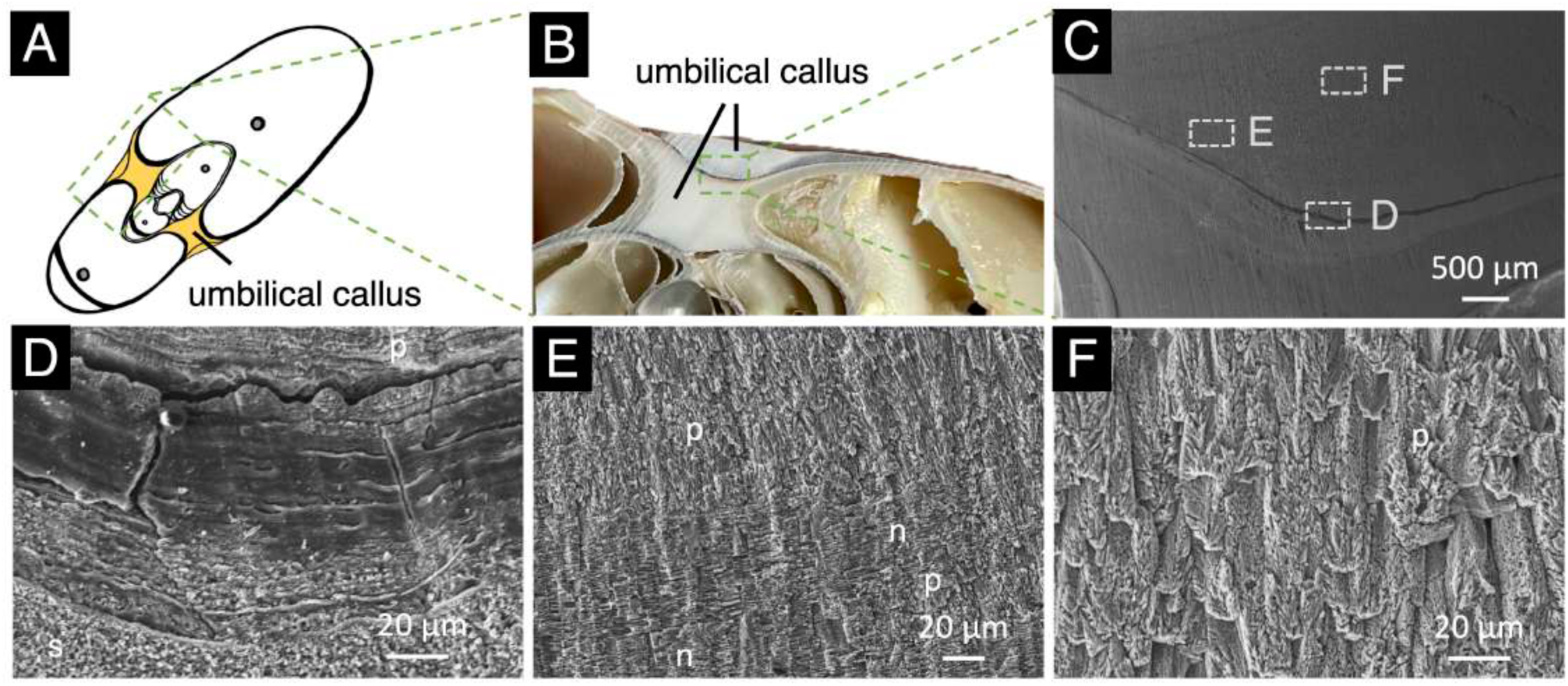
Microstructure of the umbilical callus. A) Schematic diagram of the umbilical callus. B) Photograph of the umbilical callus. C–F) SEM images showing the microstructure of the umbilical callus. D–F) Enlarged views of C. D) Organic layer, possibly corresponding to the periostracum or hood attachment site. E) Mixture of prismatic and nacreous structures in the umbilical callus. F) Enlarged view of the prismatic structure in the umbilical callus. **Abbreviations**: spherulitic structure (s), prismatic structure (p), nacreous structure (n)

**Fig. 12.**
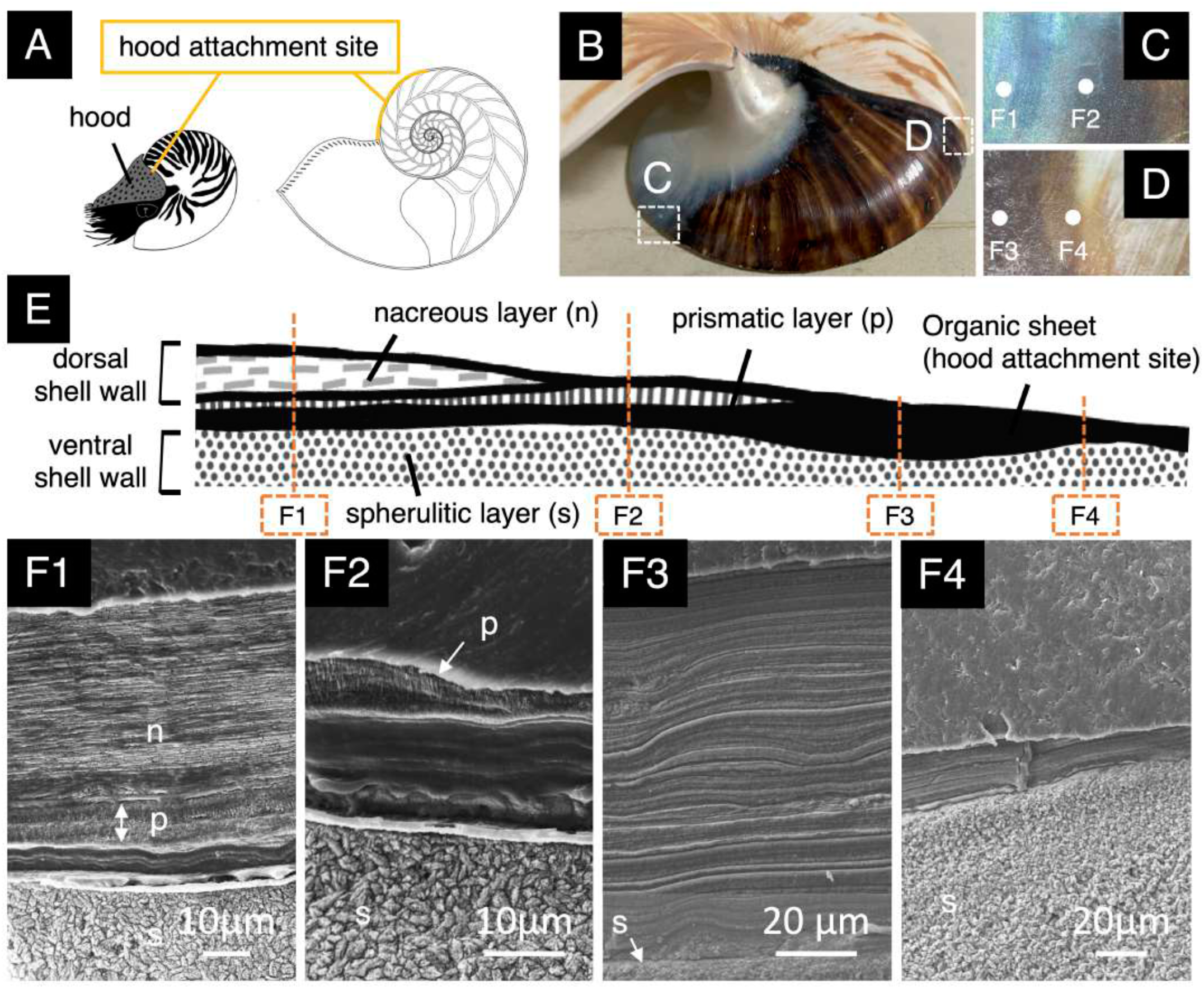
Microstructure of the hood attachment site. A) Schematic diagrams of the hood and the hood attachment site on the shell. B–D) Photographs of the hood attachment site. Enlarged surface views of B are shown in C and D. E) Schematic diagram of the microstructure around the hood attachment site. F) SEM photographs of the microstructure of the hood attachment site. F3) Thick organic layer at the hood attachment site. **Abbreviations**: spherulitic structure (s), prismatic structure (p), nacreous structure (n)

#### 5.1. Umbilical callus

The umbilical callus is a thickened area located in the depressed central region of the planispirally coiled shells (Figs. 1, 11A). In surface view, the umbilical callus exhibited black and pearly coloration extending from the dorsal shell wall (Fig. 1). Vertical sections of the umbilical callus showed crystal deposition on a thick organic layer (ca. 45 µm) derived from the previous whorl, forming a whitish basal structure that was clearly demarcated from the shell wall and septum (Fig. 11B–D). However, SEM images showed that the microstructures of the umbilical callus were largely consistent with those of the shell wall, being composed of porcellaneous and nacreous layers (Fig. 11C–F). The prismatic structure in the porcellaneous layer grew perpendicular to the outer shell surface (Fig. 11E). The prismatic structure of the umbilical callus differed slightly from that of the shell wall, showing columnar-shaped prisms rather than the triangular pyramid-shaped prisms typical of the shell wall (Fig. 11F). The nacreous layer was generally located beneath the prismatic layer, but the umbilical callus did not exhibit a simple two-layered arrangement. Instead, prismatic and nacreous structures occurred repeatedly within the umbilical callus, with gradual transitions between the two microstructures (Fig. 11E).

#### 5.2. Hood attachment site

The hood is a unique structure of the nautilus shell that function in maintaining a stable state of rest and body retraction for protection (Figs. 1, 12A). The attachment site of the hood is marked by a black deposit on the shell wall (Fig. 12B–D). Based on SEM observations, the hood attachment site consisted of laminated organic sheets and was not calcified (Fig. 12E, F). In mature specimens, the organic layer of the hood attachment site reached approximately 120 µm in thickness in regions not covered by the dorsal shell wall (Fig. 12F3). In regions sandwiched between the dorsal and ventral shell walls, the laminated organic layer was relatively thin, measuring ca. 20 µm (Fig. 12F1, 2).

## Discussion

### The *Nautilus* shell as an integrated system of biomineralization modules

The *Nautilus* shell (Fig. 1) is an integrated biomineralized structure homologous to the shell seen in other mollusks (Setiamarga et al. 2021a; Hirota et al. 2023), whose functions include body protection and buoyancy control (Denton and Gilpin-Brown 1966; Ward and Martin 1978; Greenwald et al. 1980; Greenwald et al. 1982; Greenwald and Ward 2010). However, in contrast to the shells of other mollusks, the *Nautilus* shell has distinct macrostructural components, including the shell wall, septa, and siphuncle, which are organized as an integrated system and together generate the characteristic chambered shell (Fig. 1; Table 1), suggesting that its structural logic could differ from that of other mollusks, probably in relation to selective pressures associated with the animal’s ecology, such as deep-sea habitat use (Dunstan et al. 2011a; Barord et al. 2023), and behavior, such as vertical movement and buoyancy control (Denton and Gilpin-Brown 1966; Ward and Martin 1978; Greenwald et al. 1980; Greenwald et al. 1982; Greenwald and Ward 2010), as well as hydrostatic pressure acting on the shell at depth (Kanie et al. 1980; Lemanis et al. 2016). The long evolutionary persistence of externally shelled cephalopods, which occupied dominant ecological roles in marine ecosystems from the Paleozoic to the Mesozoic (Kummel 1953; Kummel 1956; Kröger et al. 2011; Kröger 2013; Combosch et al. 2017), suggests that this shell architecture represented a functionally successful design.

At the macrostructural level, the shell wall forms the external shell surface of *Nautilus* and is comparable to the external shell of other shelled mollusks, including gastropods and bivalves, whereas the septa and siphuncle are internal structures associated with chambered cephalopods, as seen in nautiloids, ammonoids, and spirulids (Checa 2018; Lemanis et al. 2016; Checa et al. 2022; Karp et al. 2023). These components perform markedly different functions: the shell wall serves as the primary protective structure against external hazards, the septa subdivide the shell into chambers, and the siphuncle regulates buoyancy through fluid and gas exchange (Denton and Gilpin-Brown 1966; Ward and Martin 1978; Greenwald et al. 1980; Greenwald et al. 1982). Molluscan shells and their components are constructed from microscale structural units known as shell microstructures, whose arrangement affects key mechanical properties such as strength, stiffness, and toughness (e.g., Currey and Taylor 1974; Barthelat et al. 2009; Liang et al. 2016; Wan et al. 2019; Jia et al. 2022; Ghazlan et al. 2021; Deng et al. 2022; Peter et al. 2023). Molluscan shells are known to exhibit broad microstructural diversity (Carter and Clark II 1985; Carter et al. 1989; Checa 2018), probably related to their specific adaptive requirements.

Our observations show that the macrostructural complexity of the *Nautilus* shell is produced by different arrangements of five microstructural types: spherulitic, prismatic, nacreous, semi-prismatic, and irregularly oriented prismatic structures (Fig. 2). Thus, the shell wall, septum, siphuncle, myostracum, and localized shell components are not constructed from separate sets of microstructures, but from different combinations and spatial arrangements of the same five microstructural types, probably reflecting their different functional roles. Meanwhile, the presence of the same microstructural types in different shell components suggests that these components share specific structural requirements, even when their overall arrangements differ because of their different functions, indicating that certain microstructural types probably serve similar structural purposes in different macrostructural contexts (Currey and Taylor 1974; Barthelat et al. 2009; Chateigner et al. 2000; Checa 2018). For example, the nacreous structure occurs in both the shell wall and septa, probably because both components form broad load-bearing surfaces that contribute to shell strength and resistance to mechanical stress. The semi-prismatic structure and irregularly oriented prismatic structures are shared mainly by regions associated with attachment, transition, or component connection, including the inner shell wall, myostracum, septal neck, and siphuncle. Their repeated occurrence in these regions suggests that these microstructures are suited to mechanically integrate adjacent shell components, rather than to form independent shell components by themselves.

This distribution suggests that functional diversification of shell components occurred largely through reorganization of existing biomineralization modules rather than through the evolution of entirely *de novo* microstructures. Therefore, the chambered shell may represent an example of developmental modularization (Wagner et al. 2007), in which an ancestral shell-forming program became subdivided into multiple, functionally differentiated shell components. The extensive sharing of microstructural architectures among components also suggests a high degree of morphological integration (Klingenberg 2008), indicating that these modules remain developmentally and structurally connected within a common biomineralization framework.

### Microstructural transitions and mechanical integration at shell component connections

The chambered shell is essential for movement, and any structural failure, whether caused by impacts with rocky substrates or predation by durophagous organisms, is fatal (Kröger 2004; Tajika et al. 2025). Furthermore, the nautilus shell must withstand substantial changes in hydrostatic pressure resulting from repeated diel vertical migrations (Collins and Minton 1967; Dunstan et al. 2011a; Lemanis et al. 2016). Therefore, the interfaces between shell components represent critical regions where mechanical integrity must be maintained.

Mechanical performance of biominerals is shaped by the spatial organization of shell components and the arrangement and distribution of different microstructural types (Wegst et al. 2015). At the shell wall–septum interface, morphological structures such as the mural flap and adapical ridge flap, both of which are localized extensions of the shell wall that overlie the septum, have been interpreted as reinforcing elements that contribute to the mechanical strength of the chambered shell (Lemanis et al. 2020). In addition to these morphological reinforcing strategies, our observations revealed gradual transitions between different microstructural types across shell-component boundaries, suggesting that microstructural organization also contributes to mechanical integration between shell components. These transition patterns imply that the derived microstructures do not form fully discrete layers, but rather constitute a continuous construct generated by subtle modifications of a shared growth mechanism (Schoeppler et al. 2019).

Both the shell wall–septum and septum–siphuncle interfaces were characterized by gradual transitions between adjacent microstructural types. At the shell wall–septum interface, the nacreous layers of the shell wall and septum are not directly connected but are linked through an intervening semi-prismatic structure (Fig. 8). This region represents a gradual microstructural transition in which the interlamellar organic membrane characteristic of the shell-wall nacreous structure progressively disappears as the nacre grades into a semi-prismatic layer, and subsequently reappear during the transition to the septal nacreous structure. (Fig. 8G, H, J, O, P). In contrast, the septum–siphuncle interface involves a transition from nacreous to semi-prismatic structures, followed by the appearance of irregularly oriented prismatic structures adjacent to the siphuncular organic layer (Figs. 5E, 7D–F, 7H). These graded transitions may distribute stress more effectively, suppress crack propagation, and promote crack deflection, analogous to the mechanical advantages of functionally graded materials (FGMs), in which gradual structural transitions reduce stress concentrations and improve fracture resistance (Birman and Byrd 2007; Miyamoto et al. 2013; Birman 2014; Wallis et al. 2022). These observations suggest that microstructural transitions are not only a consequence of shell growth, but also constitute an important architectural strategy that facilitates both structural integration and mechanical stability in the chambered shell.

### Microstructural differentiation and functional specialization within the shell wall

The arrangement and thickness of shell microstructures are closely linked to the mechanical performance of biominerals (Liang et al. 2020). In the present study, we revealed marked dorsoventral differences in both the thickness and distribution of shell-wall microstructures (Fig. 3). The outer porcellaneous layer and inner nacreous layer progressively thickened throughout ontogeny in the ventral shell wall, but the dorsal shell wall remained relatively thin throughout ontogeny (Fig. 3D). These contrasting growth patterns suggest that shell-wall architecture is regionally differentiated according to localized functional requirements.

The ventral shell wall is associated with mechanical demands during interactions with various external environments. Increased investment in mechanically resistant microstructures within the ventral shell wall may enhance resistance to impact and abrasion, thereby reducing the risk of structural failure. The porcellaneous layer, composed of spherulitic and prismatic sub-layers, is particularly important in resisting localized damage and wear because these microstructures can dissipate deformation energy and improve resistance to abrasion and penetration (Liang et al. 2021a, b). The granule-shaped, isotropic spherulitic structure of the porcellaneous layer provides superior resistance to perforation and wear because the crystals can rotate more readily under deformation than those in prismatic or nacreous structures, dissipating energy and enhancing nanoscale plasticity (Li et al. 2006). Furthermore, the laminated, tablet-shaped structure of the inner nacreous layer confers outstanding crack resistance, as cracks propagate tortuously along soft intertablet interfaces, dissipating substantial amounts of energy (Liang et al. 2021b; Jia et al. 2022). Therefore, the progressive thickening of the ventral shell layers during ontogeny may reflect increasing mechanical demands associated with body growth, helping to maintain shell integrity under progressively greater mechanical loads.

The dorsal shell wall does not require extensive structural thickening during ontogeny. Rather than reinforcing the shell through continued deposition, dorsal shell-wall growth may be constrained by the need to maintain structural integration with previously formed whorls while preserving sufficient internal volume within the body chamber. Collectively, these patterns indicate that functional specialization can arise within a single shell component through localized regulation of shell formation.

### Early shell formation and variability of the caecum

The embryonic shell structures are shaped primarily by developmental processes that establish subsequent shell architecture, and therefore often reflect developmental constraints more strongly than the functional demands encountered later in ontogeny (e.g., Alberch 1982). Variation in these early-forming structures can provide valuable insights into the developmental flexibility of shell construction that are not evident from later-formed shell components (e.g., Klingenberg 2008). The shell formation of the nautilus proceeds sequentially throughout an embryonic developmental period of approximately ten months (Saunders 1983; Tanabe and Uchiyama 1997; Shigeno et al. 2008). Following the initiation of the protoseptum, the shell wall begins to develop, and the siphuncular tube is established only after the protoseptum has been completed (Fig. 9). The caecum, which forms the initial part of the siphuncle, was structurally variable between two specimens (Fig. 9).

Although the two specimens, *N. pompilius* and *N. cf. vitiensis*, possessed a superficially similar caecum, their microstructural organization differed markedly (Fig. 9). In *N. pompilius*, the caecum was primarily formed through a curvature of the septal nacreous layer, whereas the caecum of *N. cf. vitiensis* incorporated an additional organic layer and lacked the pronounced curvature observed in *N. pompilius*. This divergence in microstructural organization suggests that multiple constructional pathways may be capable of producing equivalent morphological traits and may indicate that the caecum is subject to weaker developmental constraints than other shell components, allowing alternative combinations of shell-forming modules to generate a functional structure while maintaining its overall morphology. A comparable pattern has been reported in *Spirula*, in which a caecum with a morphology superficially resembling the bactritoid/ammonoid type was shown to possess a distinct microstructural organization, suggesting an alternative developmental origin from the belemnoid type (Fuchs 2019). Therefore, we suggest that the caecum represents a structurally flexible region of the chambered cephalopod shell, where substantial variation in shell construction can occur without corresponding changes in overall morphology.

Alternatively, the distinct microstructures of the caecum may reflect species-specific differences or environmental influences acting during embryonic development. In this study, the two examined specimens, *N. pompilius* and *N. cf. vitiensis*, represent distant lineages within extant Nautilidae and therefore may differ in aspects of shell construction. Moreover, environmental effects during embryonic development cannot be excluded. Hatching size and embryonic developmental period in *Nautilus* exhibit substantial variation, with broad overlap among species (Tajika et al. 2022). Neither habitat depth nor ambient water temperature alone adequately explains this variation, suggesting that other factors, including water chemistry and the nutritional status of the yolk sac, may also contribute (Tajika et al. 2022). Given the exceptionally long embryonic developmental period of approximately ten months, even subtle differences in developmental conditions may influence the formation of early shell structures such as the caecum. Although the limited number of specimens examined here, reflecting the rarity and conservation concerns associated with extant nautilids, precludes robust conclusions regarding the origin of this variation, the results suggest that the caecum may represent one of the most developmentally flexible regions of the nautilus shell and may preserve evidence of alternative pathways of shell construction that are not apparent at the macroscopic level.

### The *Nautilus* shell as a modular extended phenotype

Extended phenotypes, such as beaver dams (Dawkins 2004) and spider webs (Japyassú and Laland 2017; Nakata 2012; Blamires et al. 2018), are phenotypic structures formed outside an organism’s anatomical body through genetic, developmental, physiological, or behavioral control, and their presence affects survival, reproduction, behavior, or ecological interaction (Dawkins 1982; Hunter 2009; Laland et al. 2015; Hunter 2018). The argonaut eggcase is also an example of an extended phenotype (Hirota et al. 2026). It is a non-homologous shell-like biomineralized structure secreted outside the body, used for egg protection and buoyancy control, actively formed and repaired, and interpreted as a convergently evolved extended phenotype supported by genomic evidence for shell-formation-related genes in Argonauta (Yoshida et al. 2022; Hirota et al. 2026).

Molluscan shell formation is mediated mainly by the mantle epithelium, which secretes organic matrices and shell matrix proteins that regulate calcium carbonate deposition, crystal nucleation, crystal growth, mineral polymorph selection, and shell-layer formation (Marin et al. 2013; Kocot et al. 2016; McDougall and Degnan 2018; Jackson 2021). Comparative shell-proteomic studies show that conchiferan shell formation and biomineralization use shared molecular components (Setiamarga et al. 2021a; Hirota et al. 2023), while developmental studies show that shell morphological formation is developmentally controlled, with developmental gene-expression patterns associated with shell-field patterning in mollusks (Jacobs et al. 2000; Nederbragt et al. 2002; Wollesen et al. 2017). Molluscan shells can thus be interpreted as extended phenotypes because they are not simply external mineral deposits, but external structures that affect the survival of the organism and are formed through genetically controlled processes.

In *Nautilus*, shell organization is not uniform. The shell wall, septa, siphuncle, myostracum, and localized shell components can be interpreted as developmental modules because each component is produced as a regionally differentiated part of the same shell-forming system (Wagner et al. 2007; Klingenberg 2008; Jackson 2021). These modules do not require separate microstructural systems. Instead, they are formed by changing the local order, position, and combination of shared microstructural types, so that the same microstructures act as reusable units in different parts of the shell (Chateigner et al. 2000; Checa 2018). For example, one component may be dominated by a nacreous arrangement, another may combine nacreous and prismatic structures, and another may use semi-prismatic or irregularly oriented prismatic structures around attachment, transition, or connection zones. Although the final shell is continuous, its component-level differences are produced by using the same microstructural units in different arrangements within a single organism. This organization represents developmental modularity, because distinct shell components are generated through regional rearrangement of shared structural elements rather than through independent construction of each component (Wagner et al. 2007; Klingenberg 2008; Jackson 2021). The underlying genetic components of shell formation may similarly be rearranged or redeployed in different regional combinations to produce distinct microstructures, and these microstructures are then combined into larger shell components (True and Carroll 2002; Herlitze et al. 2018; Jackson 2021). The *Nautilus* shell can therefore be interpreted as an extended phenotype with modular organization, in which genetically controlled shell formation produces an external structure subdivided into distinct shell components while maintaining continuity through shared microstructural construction and gradual transitions across component boundaries (Dawkins 1982; Dawkins 2004; Wagner et al. 2007; Klingenberg 2008; Jackson 2021).

### Evolutionary implications of shell microstructural organization in cephalopods

The present study provides a comprehensive characterization of the diversity, distribution, and organization of shell microstructures throughout the chambered shell of *Nautilus*. As the only extant cephalopod lineage retaining an external chambered shell, *Nautilus* occupies a unique position for understanding the construction and functional significance of ancestral externally shelled cephalopod architectures (e.g., Webers and Yochelson 1989; Sasaki et al. 2010; Shigeno et al. 2010; Ritterbush et al. 2014; Pohle et al. 2022). The nautiloid lineage, including extant nautilus, originated from a monoplacophoran-like conchiferan in the Late Cambrian (Kröger et al. 2011; Setiamarga 2021a, b), and many aspects of its shell organization are thought to retain features of the ancestral cephalopods. Consequently, the microstructural architecture documented here provides valuable insights into the organization and functional roles of shell characters that were widespread among extinct externally shelled cephalopods.

Although microstructural data from most extinct nautiloid groups remain limited because of preservation constraints, extant nautilids retain several ancestral shell features that permit meaningful comparisons with phylogenetically distinct chambered cephalopods, including ammonites, belemnites, spirulids, and cuttlefish (Tanabe et al. 1982; Checa et al. 2015; Checa et al. 2022). The repeated deployment of a limited set of microstructural types across multiple shell components, together with their integration through gradual transitions at component boundaries, suggests that the complexification of the cephalopod shell may have proceeded largely through the reorganization and specialization of existing biomineralization modules. The shell microstructural architecture of *Nautilus* provides a basis for comparing evolutionary transformations in cephalopods, from ancestral chambered-shell architectures to lineage-specific modifications. Our integrative morphomics analysis links microstructural diversity, spatial organization, ontogenetic variation, and structural connections across functional components, showing that the *Nautilus* shell is an integrated system of biomineralization modules.

## Acknowledgments

DHES and KH thank past and present members of the Setiamarga Lab at NITW and Sasaki Lab at The University of Tokyo for their support throughout this study. DHES thanks Oleg Simakov (University of Vienna) and Linda and Nick Holland (Scripps Institution of Oceanography) for their support during DHES’ sabbatical stay at University of Vienna.

## Author contributions

KH, TS, and DHES conceived the study. TS and DHES supervised the project. KH conducted the main microstructural analyses. KH and TS collected the samples and associated specimen information. KH wrote the first draft of the manuscript and KH, TS, and DHES wrote subsequent versions of the manuscript. All authors were involved in data interpretation and discussions. All authors read, edited, and confirmed the content of the final version of the manuscript.

## Data availability

All specimens examined in this study are deposited in The University Museum, The University of Tokyo. All data supporting the findings of this study are available in the manuscript, and additional information is provided in the Supplementary Figures.

## Supplementary Figures

**Fig. S1.** Identification of nautilus species based on morphological characteristics. Photographs showing the entire shell and pigmented shell regions of two specimens: A) *N. pompilius* and B) *N. cf. vitiensis*. C) Characterization map of *Nautilus* shell morphologies (Barord et al. 2023).

**Fig. S2.** Microstructure of the shell wall during early ontogeny. Microstructure of the shell wall during early shell development. A) Schematic diagram of the shell at an early developmental stage. SEM images showing microstructures on the 1) ventral and 2) dorsal sides. Abbreviations: spherulitic and prismatic structure (s-p); prismatic structure (p); nacreous structure (n).

## References

1. Alberch P (1982) Developmental constraints in evolutionary processes. In Evolution and Development: Report of the Dahlem Workshop on Evolution and Development Berlin 1981, May 10–15. Berlin, Heidelberg: Springer Berlin Heidelberg. 313–332. 10.1007/978-3-642-45532-2_15

2. Arnold JM, Carlson BA (1986) Living *Nautilus* embryos: preliminary observations. Science 232:73–76. 10.1126/science.232.4746.73

3. Arnold JM (2010) Reproduction and embryology of Nautilus. In Nautilus: The Biology and Paleobiology of a Living Fossil, Reprint with additions. Dordrecht: Springer Netherlands. 353–372. 10.1007/978-90-481-3299-7_26

4. Bandel K, Boletzky SV (1979) A comparative study of the structure, development and morphological relationships of chambered cephalopod shells. Veliger 21:313–354. https://oceanrep.geomar.de/id/eprint/38116

5. Bandel K (1989) Cephalopod shell structure and general mechanisms of shell formation. Skeletal biomineralization: patterns, processes and evolutionary trends 5:97–115. 10.1029/SC005p0097

6. Barord GJ, Dooley F, Dunstan A, Ilano A, Keister KN, Neumeister H, Preuss T, Schoepfer S, Ward PD (2014) Comparative population assessments of Nautilus sp. in the Philippines, Australia, Fiji, and American Samoa using baited remote underwater video systems. PLoS One 9:e100799. 10.1371/journal.pone.0100799

7. Barord GJ, Beydoun M, Bruce S, Li V, Ward PD, Basil J (2021) Foraging and scavenging in nautilus (Nautilus sp.) L. (Cl. Cephalopoda). Mar Freshw Behav Physiol 54:241–261. 10.1080/10236244.2021.2003195

8. Barthelat F, Rim JE, Espinosa HD. (2009) A review on the structure and mechanical properties of mollusk shells–perspectives on synthetic biomimetic materials. Applied Scanning Probe Methods XIII: Biomimetics And Industrial Applications 17–44. 10.1007/978-3-540-85049-6_2

9. Barord GJ, Combosch DJ, Giribet G, Landman N, Lemer S, Veloso J, Ward PD (2023) Three new species of Nautilus Linnaeus, 1758 (Mollusca, Cephalopoda) from the Coral Sea and South Pacific. ZooKeys 1143:51–69. 10.3897/zookeys.1143.84427

10. Basil JA, Hanlon RT, Sheikh SI, Atema J (2000) Three-dimensional odor tracking by *Nautilus pompilius*. J Exp Biol 203:1409–1414. 10.1242/jeb.203.9.1409

11. Basil J, Bahctinova I, Kuroiwa K, Lee N, Mims D, Preis M, Soucier C (2005) The function of the rhinophore and the tentacles of *Nautilus pompilius* L. (Cephalopoda, Nautiloidea) in orientation to odor. Mar Freshw Behav Physiol 38:209–221. 10.1080/10236240500310096

12. Birman V, Byrd LW (2007) Modeling and analysis of functionally graded materials and structures. Appl Mech Rev 60:195–216. 10.1115/1.2777164

13. Birman V (2014) Functionally Graded Materials and Structures. In: Hetnarski, R.B. (eds) Encyclopedia of Thermal Stresses. Springer, Dordrecht. 10.1007/978-94-007-2739-7_573

14. Blamires SJ, Martens PJ, Kasumovic MM. (2018) Fitness consequences of plasticity in an extended phenotype. J Exp Biol 221(4):jeb167288. 10.1242/jeb.167288

15. Boutilier RG, West TG, Webber DM, Pogson GH, Mesa KA, Wells J, Wells MJ (2000) The protective effects of hypoxia-induced hypometabolism in the *Nautilus*. J Comp Physiol B 170:261–268. 10.1007/s003600000096

16. Chateigner D, Hedegaard C, Wenk HR (2000) Mollusc shell microstructures and crystallographic textures. J Struct Geol 22(11–12):1723–1735. 10.1016/S0191-8141(00)00088-2

17. Carter JG, Clark II GR (1985) Classification and phylogenetic significance of molluscan shell microstructure. Series in Geology, Notes for Short Course 13:50–71. 10.1017/S0271164800001093

18. Caner JG, Bandel K, de Buffrénil V, Carlson S, Castanet J, Dalingwater J, Francillon-Vieillot H, Géraudie J, Meunier FJ, Mutvei H, de Ricqlès A, Sire JY, Smith A, Wendt J, Williams A, Zylberberg L (1989) Glossary of skeletal biomineralization. Skeletal Biomineralization: Patterns, Processes and Evolutionary Trends 5:337–352. 10.1002/9781118667279.oth1

19. Checa AG, Cartwright JH, Sánchez-Almazo I, Andrade JP, Ruiz-Raya F (2015) The cuttlefish *Sepia officinalis* (Sepiidae, Cephalopoda) constructs cuttlebone from a liquid-crystal precursor. Sci Rep 5(1):11513. 10.1038/srep11513

20. Checa AG (2018) Physical and biological determinants of the fabrication of molluscan shell microstructures. Front Mar Sci 5:353. 10.3389/fmars.2018.00353

21. Checa AG, Grenier C, Griesshaber E, Schmahl WW, Cartwright JH, Salas C, Oudot M (2022) The shell structure and chamber production cycle of the cephalopod *Spirula* (Coleoidea, Decabrachia). Mar Biol 169:132. 10.1007/s00227-022-04120-0

22. Clark MS, Peck LS, Arivalagan J, Backeljau T, Berland S, Cardoso JC, Caurcel C, Chapelle G, De Noia M, Dupont S, Gharbi K, Hoffman JI, Last KS, Marie A, Melzner F, Michalek K, Morris J, Power DM, Ramesh K, Sanders T, Sillanpää K, Sleight VA, Stewart-Sinclair PJ, Sundell K, Telesca L, Vendrami DLJ, Ventura A, Wilding TA, Yarra T, Harper EM (2020) Deciphering mollusc shell production: the roles of genetic mechanisms through to ecology, aquaculture and biomimetics. Biol Rev 95:1812–1837. 10.1111/brv.12640

23. Collins DH, Minton P (1967) Siphuncular tube of *Nautilus*. Nature 216:916–917. 10.1038/216916b0

24. Collins D, Ward PD (2010) Adolescent growth and maturity in Nautilus. In Nautilus: The Biology and Paleobiology of a Living Fossil, Reprint with additions. Dordrecht: Springer Netherlands. 421–432. 10.1007/978-90-481-3299-7_29

25. Combosch DJ, Lemer S, Ward PD, Landman NH, Giribet G (2017) Genomic signatures of evolution in *Nautilus*—an endangered living fossil. Mol Ecol 26:5923–5938. 10.1111/mec.14344

26. Crick RE (1981) Diversity and evolutionary rates of Cambro-Ordovician nautiloids. Paleobiology 7:216–229. 10.1017/S0094837300003997

27. Currey JD, Taylor JD (1974) The mechanical behaviour of some molluscan hard tissues. J Zool 173(3):395–406. 10.1111/j.1469-7998.1974.tb04122.x

28. Dawkins R (1982) The extended phenotype, Oxford University Press Oxford.

29. Dawkins R (2004) Extended phenotype–but not too extended. A reply to Laland, Turner and Jablonka. Biol Philos 19(3):377–396. 10.1023/B:BIPH.0000036180.14904.96

30. Denton EJ, Gilpin-Brown JB (1966) On the buoyancy of the pearly *Nautilus*. Journal of the Marine Biological Association of the United Kingdom 46:723–759. 10.1017/S0025315400033440

31. Drushchits VV (1977) The structure of the ammonitella and the direct development of ammonites. Paleontol J 11:188.

32. Dauphin Y, Luquet G, Percot A, Bonnaud-Ponticelli L (2020) Comparison of embryonic and adult shells of *Sepia officinalis* (Cephalopoda, Mollusca). Zoomorphology 139:151–169. 10.1007/s00435-020-00477-2

33. Deng Z, Jia Z, Li L (2022) Biomineralized materials as model systems for structural composites: Intracrystalline structural features and their strengthening and toughening mechanisms. Adv Sci 9:2103524. 10.1002/advs.202103524

34. Dong W, Huang J, Liu C, Wang H, Zhang G, Xie L, Zhang R (2022) Characterization of the myostracum layers in molluscs reveals a conservative shell structure. Front Mar Sci 9:862929. 10.3389/fmars.2022.862929

35. Dunstan AJ, Ward PD, Marshall NJ (2011a) Vertical distribution and migration patterns of Nautilus pompilius. PLoS One 6:e16311. 10.1371/journal.pone.0016311

36. Dunstan AJ, Ward PD, Marshall NJ (2011b) Nautilus pompilius life history and demographics at the Osprey Reef Seamount, Coral Sea, Australia. PLoS One 6:e16312. 10.1371/journal.pone.0016312

37. Evans DH, King AH, Histon K, Cichowolski M (2014) Nautiloid cephalopods–a review of their use and potential in biostratigraphy Denisia 32:7–22. https://www.zobodat.at/pdf/DENISIA_0032_0007-0022.pdf

38. Fischer AG, Teichert C (1969) Cameral deposits in cephalopod shells. Kans Paleontol Contrib 37:1–30

39. Frey RC (1995) Middle and Upper Ordovician nautiloid cephalopods of the Cincinnati Arch region of Kentucky, Indiana, and Ohio. US Government Printing Office, Washington D.C., United States. (No. 1066). https://pubs.usgs.gov/pp/1066p/report.pdf

40. Fuchigami T, Sasaki T (2005) The shell structure of the recent Patellogastropoda (Mollusca: Gastropoda). Paleontol Res 9:143–168. 10.2517/prpsj.9.143

41. Fuchs D (2019) Homology problems in cephalopod morphology: deceptive (dis) similarities between different types of ‘caecum’. Swiss J Palaeontol 138:49–63. 10.1007/s13358-019-00183-7

42. Ghazlan A, Ngo T, Tan P, Xie, YM, Tran P, Donough M (2021) Inspiration from Nature’s body armours–A review of biological and bioinspired composites. Compos B: Eng 205:108513. 10.1016/j.compositesb.2020.108513

43. Greenwald L, Ward PD, Greenwald OE (1980) Cameral liquid transport and buoyancy control in chambered nautilus (*Nautilus macromphalus*). Nature 286:55–56. 10.1038/286055a0

44. Greenwald L, Cook CB, Ward PD (1982) The structure of the chambered *Nautilus* siphuncle: The siphuncular epithelium. J Morphol 172:5–22. 10.1002/jmor.1051720103

45. Greenwald L, Ward PD (2010) Buoyancy in *Nautilus*. In *Nautilus*: The Biology and Paleobiology of a Living Fossil, Reprint with additions. Dordrecht, Springer Netherlands. 547–560. 10.1007/978-90-481-3299-7_34

46. Grégoire C (1962) On submicroscopic structure of the Nautilus shell. Bulletin. Institut Royal des Sciences Naturelles de Belgique. Mededelingen. Koninklijk Belgisch Instituut voor Natuurwetenschappen

47. Hirano H, Obata I (1979) Shell Morphology of *Nautilus pompilius* and *N. macromphalas*. Bull Natl Mus Nat Sci, Ser C, Geol Paleontol 5:113–130.

48. Hirota K, Tochino N, Seto M, Sasaki T, Yoshida MA, Setiamarga DH (2023) Comparative proteomics of the shell matrix proteins of *Nautilus pompilius* and the conchiferans provide insights into mollusk shell evolution at the molecular level. Mar Biol 170:106. 10.1007/s00227-023-04244-x

49. Hirota K, Sasaki T, Yoshimura T, Onodera S, Hirano H, Toyama T, Yoshida MA, Setiamarga DH (2026) Microstructural insights into the functional morphology and formation logic of spherulitic–fibrous prismatic architecture in the shell–like eggcase of the argonaut octopods. Sci Rep 16(1):12372. 10.1038/s41598-026-45670-3

50. Herlitze I, Marie B, Marin F, Jackson DJ (2018) Molecular modularity and asymmetry of the molluscan mantle revealed by a gene expression atlas. GigaScience 7(6):giy056. 10.1093/gigascience/giy056

51. Hunter P (2009) Extended phenotype redux. How far can the reach of genes extend in manipulating the environment of an organism?. EMBO Rep 10(3):212. 10.1038/embor.2009.18

52. Hunter P (2018) The revival of the extended phenotype: After more than 30 years, Dawkins’ extended phenotype hypothesis is enriching evolutionary biology and inspiring potential applications. EMBO Rep 19(7):EMBR201846477. 10.15252/embr.201846477

53. Jacobs DK, Wray CG, Wedeen CJ, Kostriken R, DeSalle R, Staton JL, Gates RD, Lindberg DR (2000) Molluscan engrailed expression, serial organization, and shell evolution. Evol Dev 2(6):340–347. 10.1046/j.1525-142x.2000.00077.x

54. Jackson DJ (2021) Mantle modularity underlies the plasticity of the molluscan Shell: supporting data from *Cepaea nemoralis*. Front Genet 12:622400. 10.3389/fgene.2021.622400

55. Japyassú HF, Laland KN (2017) Extended spider cognition. Anim Cogn 20(3):375–395. 10.1007/s10071-017-1069-7

56. Jia Z, Deng Z, Li L (2022) Biomineralized materials as model systems for structural composites: 3D architecture. Adv Mater 34:2106259. 10.1002/adma.202106259

57. Kanie Y, Fukuda Y, Nakayama H, Seki K, Hattori M (1980) Implosion of living *Nautilus* under increased pressure. Paleobiology 6:44–47. 10.1017/S0094837300012483

58. Karp MA, Phillips B, Edie SM (2023) Investigations into 3D-printed nautiloid-inspired pressure housings. Bioinspir Biomim 18:066015. 10.1088/1748-3190/acfeb8

59. Kiel S, Goedert JL, Tsai CH (2022) Seals, whales and the Cenozoic decline of nautiloid cephalopods. J Biogeogr 49:1903–1910. 10.1111/jbi.14488

60. Klingenberg CP (2008) Morphological integration and developmental modularity. Annu Rev Ecol Syst 39(1):115–132. 10.1146/annurev.ecolsys.37.091305.110054

61. Kocot KM, Aguilera F, McDougall C, Jackson DJ, Degnan BM (2016) Sea shell diversity and rapidly evolving secretomes: insights into the evolution of biomineralization. Front Zool 13(1):23. 10.1186/s12983-016-0155-z

62. Kummel B (1953) American Triassic coiled nautiloids (No. 250). Profess Pap U.S. Geol Surv 250:1–104 10.3133/pp250

63. Kummel B (1956) Post-Triassic nautiloid genera. Bull Mus Comp Zool 114:324–494.

64. Kröger B (2004) Large shell injuries in Middle Ordovician Orthocerida (Nautiloidea, Cephalopoda). GFF 126(3):311–316. 10.1080/11035890401263311

65. Kröger B, Vinther J, Fuchs D (2011) Cephalopod origin and evolution: a congruent picture emerging from fossils, development and molecules: extant cephalopods are younger than previously realised and were under major selection to become agile, shell-less predators. BioEssays 33:602–613. 10.1002/bies.201100001

66. Kröger B (2013) The cephalopods of the Boda Limestone, Late Ordovician, of Dalarna, Sweden. Eur J Taxon 41. 10.5852/ejt.2013.41

67. Laland KN, Uller T, Feldman MW, Sterelny K, Müller GB, Moczek A, Jablonka E, Odling-Smee J (2015) The extended evolutionary synthesis: its structure, assumptions and predictions. Proc R Soc B: Biol Sci 282(1813):20151019. 10.1098/rspb.2015.1019

68. Landman NH, Arnold JM, Mutvei H (1989) Description of the embryonic shell of *Nautilus belauensis* (Cephalopoda). Am Mus Novit 2960:1–16

69. Landman NH, Cochran JK (2010) Growth and longevity of *Nautilus*. In *Nautilus*: The Biology and Paleobiology of a Living Fossil, Reprint with additions. Dordrecht, Springer Netherlands. 401–420. 10.1007/978-90-481-3299-7_28

70. Lee SW, Jang YN, Kim JC (2011) Characteristics of the aragonitic layer in adult oyster shells, Crassostrea gigas: structural study of myostracum including the adductor muscle scar. Evid Based Complement Alternat Med 2011:742963. 10.1155/2011/742963

71. Lemanis R, Zachow S, Hoffmann R (2016) Comparative cephalopod shell strength and the role of septum morphology on stress distribution. PeerJ 4:e2434. 10.7717/peerj.2434

72. Lemanis R, Stier D, Zlotnikov I, Zaslansky P, Fuchs D (2020) The role of mural mechanics on cephalopod palaeoecology. J. R. Soc. Interface 17:20200009. 10.1098/rsif.2020.0009

73. Liang Y, Zhao Q, Li X, Zhang Z, Ren L (2016) Study of the microstructure and mechanical properties of white clam shell. Micron 87:10–17. 10.1016/j.micron.2016.04.007

74. Liang SM, Ji HM, Li XW (2020) Thickness-dependent mechanical properties of nacre in *Cristaria plicata* shell: Critical role of interfaces. J Mater Sci Technol 44:1–8. 10.1016/j.jmst.2019.10.039

75. Liang SM, Ji HM, Li XW (2021a) A high-strength and high-toughness nacreous structure in a deep-sea *Nautilus* shell: Critical role of platelet geometry and organic matrix. J Mater Res Technol 88:189–202. 10.1016/j.jmst.2021.01.082

76. Liang SM, Ji HM, Li YY, Li XW (2021b) An ingenious microstructure arrangement in deep-sea Nautilus shell against the harsh environment. ACS Biomater Sci Eng 7:4819–4827. https://orcid.org/0000-0002-0238-9107

77. Li X, Xu ZH, Wang R (2006) *In situ* observation of nanograin rotation and deformation in nacre. Nano Lett 6:2301–2304. 10.1021/nl061775u

78. Lowenstam HA, Traub W, Weiner S (1984) *Nautilus* hard parts: a study of the mineral and organic constitutents. Paleobiology 10:268–279. 10.1017/S0094837300008198

79. Marin F, Marie B, Hamada SB, Ramos-Silva P, Le Roy N, Guichard N, Wolf S, Montagnani C, Joubert C, Piquemal D, Saulnier D, Gueguen Y (2013) Shellome’: Proteins involved in mollusk shell biomineralization-diversity, functions. Recent Adv Pearl Res 33:149–166.

80. McDougall C, Degnan BM (2018) The evolution of mollusc shells. Wiley Interdiscip Rev Dev Biol 7(3):e313. 10.1002/wdev.313

81. Miyamoto Y, Kaysser WA, Rabin BH, Kawasaki A, Ford RG (2013) (Eds.) Functionally graded materials: design, processing and applications. Springer Science & Business Media.

82. Mutvei H (1964) On the shells of Nautilus and Spirula with notes on the shell secretion in non-cephalopod mollusks. Ark. Zool 16:221–278.

83. Mutvei H (1972) Ultrastructural studies on cephalopod shells. I. The septa and siphonal tube in Nautilus. Bull Geol Inst Univ Uppsala 3:237–261.

84. Mutvei H, Doguzhaeva L (1997) Shell ultrastructure and ontogenetic growth in *Nautilus pompilius* L.(Mollusca: Cephalopoda). Palaeontogr Abt A: Palaozool-Stratigr 246:33–52. 10.1127/pala/246/1997/33

85. Mutvei H (2017) Siphuncular structure in the extant *Spirula* and in other coleoids (Cephalopoda). GFF 139:129–139. 10.1080/11035897.2016.1227364

86. Nakata K (2012) Plasticity in an extended phenotype and reversed up-down asymmetry of spider orb webs. Anim Behav 83(3):821–826. 10.1016/j.anbehav.2011.12.030

87. Nederbragt AJ, van Loon AE, Dictus WJ (2002) Expression of *Patella vulgata* orthologs of engrailed and dpp-BMP2/4 in adjacent domains during molluscan shell development suggests a conserved compartment boundary mechanism. Dev Biol 246(2):341–355. 10.1006/dbio.2002.0653

88. Owen R. Memoir on the pearly *Nautilus* (*Nautilus pompilius*, linn.). Richard Taylor, London, UK. 1832.

89. Peter NJ, Griesshaber E, Reisecker C, Hild S, Oliveira MV, Schmahl WW, Schneider AS (2023) Biocrystal assembly patterns, biopolymer distribution and material property relationships in *Mytilus galloprovincialis*, Bivalvia, and *Haliotis glabra*, Gastropoda, shells. Materialia 28:101749. 10.1016/j.mtla.2023.101749

90. Pietsch C, Anderson BM, Maistros LM, Padalino EC, Allmon WD (2021) Convergence, parallelism, and function of extreme parietal callus in diverse groups of Cenozoic Gastropoda. Paleobiology 47:337–362. 10.1017/pab.2020.33

91. Pohle A, Kröger B, Warnock RC, King AH, Evans DH, Aubrechtová M, Cichowolski M, Fang X, Klug C (2022) Early cephalopod evolution clarified through Bayesian phylogenetic inference. BMC Biol 20:88. 10.1186/s12915-022-01284-5

92. Pohle A, Hoffmann R, Nützel A, Seuss B, Aubrechtová M, Kröger B, Stevens K, Immenhauser A (2025) Microstructural and geochemical evidence offers a solution to the cephalopod cameral deposits riddle. Palaeontology 68(6):e70032. 10.1111/pala.70032

93. Ritterbush KA, Hoffmann R, Lukeneder A, De Baets K (2014) Pelagic palaeoecology: the importance of recent constraints on ammonoid palaeobiology and life history. J Zool 292:229–241. 10.1111/jzo.12118

94. Saunders WB (1983) Natural rates of growth and longevity of *Nautilus belauensis*. Paleobiology 9:280–288. 10.1017/S0094837300007697

95. Saunders WB (1984a) The role and status of *Nautilus* in its natural habitat: evidence from deep-water remote camera photosequences. Paleobiology 10:469–486. 10.1017/S0094837300008472

96. Saunders WB (1984b) *Nautilus* growth and longevity: evidence from marked and recaptured animals. Science 224:990–992. 10.1126/science.224.4652.990

97. Saunders WB, Landman N (Eds.) 2009 Nautilus: the biology and paleobiology of a living fossil, reprint with additions Springer Science & Business Media, Berlin, Germany. 10.1007/978-90-481-3299-7

98. Sasaki T, Shigeno S, Tanabe K, Shigeta Y, Hirano H (2010) Anatomy of living Nautilus: Reevaluation of primitiveness and comparison with Coleoidea. Cephalopods—Present and past, Tokai University Press, Tokyo 35–66.

99. Sato K, Sasaki T (2015) Shell microstructure of Protobranchia (Mollusca: Bivalvia): diversity, new microstructures and systematic implications. Malacologia 59:45–103. 10.4002/040.059.0106

100. Schneider CA, Rasband WS, Eliceiri KW (2012) NIH Image to ImageJ: 25 years of image analysis. Nat Methods 9(7):671–675. 10.1038/nmeth.2089

101. Schoeppler V, Lemanis R, Reich E, Pusztai T, Gránásy L, Zlotnikov I (2019) Crystal growth kinetics as an architectural constraint on the evolution of molluscan shells. Proc Natl Acad Sci U S A 116:20388–20397. 10.1073/pnas.1907229116

102. Schöne BR, Dunca E, Fiebig J, Pfeiffer M (2005) Mutvei’s solution: an ideal agent for resolving microgrowth structures of biogenic carbonates. Palaeogeogr Palaeoclimatol Palaeoecol 228(1–2):149–166. 10.1016/j.palaeo.2005.03.054

103. Setiamarga DH, Hirota K, Yoshida MA, Takeda Y, Kito K, Ishikawa M, Shimizu K, Isowa Y, Ikeo K, Sasaki T, Endo K (2021a) Hydrophilic shell matrix proteins of *Nautilus pompilius* and the identification of a core set of conchiferan domains. Genes 12:1925. 10.3390/genes12121925

104. Setiamarga DH (2021b) Evolutionary saga of the cephalopod shell: A molecular paleontological approach. Aquabiology 43(3):270–279 (In Japanese with English abstract)

105. Son R, Yamazawa K, Oguchi A, Suga M, Tamura M, Yanagita M, Murakawa Y, Kume S (2023) Morphomics via next-generation electron microscopy. J Mol Cell Biol 15(12):mjad081. 10.1093/jmcb/mjad081

106. Shigeno S, Sasaki T, Moritaki T, Kasugai T, Vecchione M, Agata K (2008) Evolution of the cephalopod head complex by assembly of multiple molluscan body parts: evidence from *Nautilus* embryonic development. J. Morphol 269:1–17. 10.1002/jmor.10564

107. Shigeno S, Sasaki T, Boletzky SV (2010) The origins of cephalopod body plans: a geometrical and developmental basis for the evolution of vertebrate-like organ systems. Cephalopods-Present and Past 1:23–34.

108. Tajika A, Landman NH, Slovacek M, Nishida K, Morita W, Witts JD (2022) Intra-and interspecific variability in offspring size in nautilids. Lethaia 55(3):1–17. 10.18261/let.55.3.1

109. Tajika A, Landman NH, Cochran JK et al. Ammonoid extinction versus nautiloid survival: is metabolism responsible?. Geology 2023;51:621–625. 10.1130/G51116.1

110. Tajika A, Rashkova A, Landman NH, Klompmaker AA (2025) Lethal injuries on the scaphitid ammonoid Hoploscaphites nicolletii (Morton, 1842) in the Upper Cretaceous Fox Hills Formation, South Dakota, USA. Swiss J Palaeontol 144(1):1. 10.1186/s13358-024-00341-6

111. Tanabe K, Fukuda Y, Obata I (1982) Formation and function of the siphuncle-septal neck structures in two Mesozoic ammonites. Trans Proc Palaeont Soc Japan, N.S. 128:433–443. 10.14825/prpsj1951.1982.128_433

112. Tanabe K, Uchiyama K (1997) Development of the embryonic shell structure in Nautilus. Veliger 40:203–215.

113. Teichert C, Matsumoto T (2010) The ancestry of the genus *Nautilus*. In *Nautilus*: the biology and paleobiology of a living fossil, reprint with additions. Dordrecht, Springer Netherlands 25–32. 10.1007/978-90-481-3299-7_2

114. Thomas RDK (1988) Evolutionary convergence of bivalved shells: a comparative analysis of constructional constraints on their morphology. Am Zool 28:267–276. 10.1093/icb/28.1.267

115. True JR, Carroll SB (2002) Gene co-option in physiological and morphological evolution. Annu Rev Cell Dev Biol 18(1):53–80. 10.1146/annurev.cellbio.18.020402.140619

116. Tanner A R, Fuchs D, Winkelmann IE, Gilbert MTP, Pankey MS, Ribeiro ÂM, Kocot KM, Halanych KM, Oakley TH, da Fonseca RR, Pisani D, Vinther J (2017) Molecular clocks indicate turnover and diversification of modern coleoid cephalopods during the Mesozoic Marine Revolution. Proc R Soc B: Biol Sci 284(1850):20162818. 10.1098/rspb.2016.2818

117. Vermeij GJ (1988) The Natural History of *Nautilus*. Science 240:1074–1075. 10.1126/science.240.4855.1074

118. Vinn O, ten Hove HA (2011) Microstructure and formation of the calcareous operculum in Pyrgopolon ctenactis and Spirobranchus giganteus (Annelida, Serpulidae). Zoomorphology 130:181–188. 10.1007/s00435-011-0133-0

119. Vinn O (2013) On the unique isotropic aragonitic tube microstructure of some serpulids (Polychaeta, Annelida). J Morphol 274:478–482. 10.1002/jmor.20112

120. Wagner GP, Pavlicev M, Cheverud JM (2007) The road to modularity. Nat Rev Genet 8(12):921–931. 10.1038/nrg2267

121. Walker SE, Brett CE (2002) Post-Paleozoic patterns in marine predation: was there a Mesozoic and Cenozoic marine predatory revolution?. Paleontol Soc Pap 8:119–194. 10.1017/S108933260000108X

122. Wallis D, Harris J, Böhm CF, Wang D, Zavattieri P, Feldner P, Merle B, Pipich V, Hurle K, Leupold S, Hansen LN, Marin F, Wolf SE (2022) Progressive changes in crystallographic textures of biominerals generate functionally graded ceramics. Mater Adv 3(3):1527–1538. 10.1039/d1ma01031j

123. Wan C, Ma Y, Gorb SN (2019) Compromise between mechanical and chemical protection mechanisms in the *Mytilus edulis* shell. J Exp Biol 222:jeb201103. 10.1242/jeb.201103

124. Ward P, Martin AW (1978) On the buoyancy of the pearly *Nautilus*. J Exp Zool 205:5–12. 10.1002/jez.1402050103

125. Ward P, Greenwald L, Magnier Y (1981) The chamber formation cycle in *Nautilus macromphalus*. Paleobiology 7:481–493. 10.1017/S0094837300025537

126. Ward P, Dooley F, Barord GJ (2016) *Nautilus*: biology, systematics, and paleobiology as viewed from 2015. Swiss J Palaeontol 135:169–185. 10.1007/s13358-016-0112-7

127. Webers GF, Yochelson EL (1989) Late Cambrian molluscan faunas and the origin of the Cephalopoda. Geol Soc Spec Publ 47:29–42. 10.1144/GSL.SP.1989.047.01.0

128. Wegst UG, Bai H, Saiz E, Tomsia AP, Ritchie RO (2015) Bioinspired structural materials. Nat Mater 14(1):23–36. 10.1038/nmat4089

129. Wells MJ, Wells J (1985) Ventilation and oxygen uptake by *Nautilus*. J Exp Biol 118:297–312. 10.1242/jeb.118.1.297

130. Westermann GE (1973) Strength of concave septa and depth limits of fossil cephalopods. Lethaia 6:383–403. 10.1111/j.1502-3931.1973.tb01205.x

131. Westermann B, Beck-Schildwächter I, Beuerlein K, Kaleta EF, Schipp R (2004) Shell growth and chamber formation of aquarium-reared Nautilus pompilius (Mollusca, Cephalopoda) by X-ray analysis. J Exp Zool A Comp Exp Biol 301:930–937. 10.1002/jez.a.116

132. Westermann B, Beuerlein K (2005) Y-maze experiments on the chemotactic behaviour of the tetrabranchiate cephalopod *Nautilus pompilius* (Mollusca). Mar Biol 147:145–151. 10.1007/s00227-005-1555-3

133. Wollesen T, Scherholz M, Rodriguez Monje SV, Redl E, Todt C, Wanninger A (2017) Brain regionalization genes are co-opted into shell field patterning in Mollusca. Sci Rep 7(1):5486. 10.1038/s41598-017-05605-5

134. Wu S, Chiang CY, Zhou W (2017) Formation mechanism of CaCO3 spherulites in the myostracum layer of limpet shells. Crystals 7:319. 10.3390/cryst7100319

135. Yoshida MA, Hirota K, Imoto J, Okuno M, Tanaka H, Kajitani R, Toyoda A, Itoh T, Ikeo K, Sasaki T, Setiamarga DH (2022) Gene recruitments and dismissals in the argonaut genome provide insights into pelagic lifestyle adaptation and shell-like eggcase reacquisition. GBE 14(11): evac140. 10.1093/gbe/evac140

