## Supplementary Files for "Nautilus shell morphomics reveals microstructural heterogeneity alongside structural continuity across component boundaries"

### Supplementary Figures

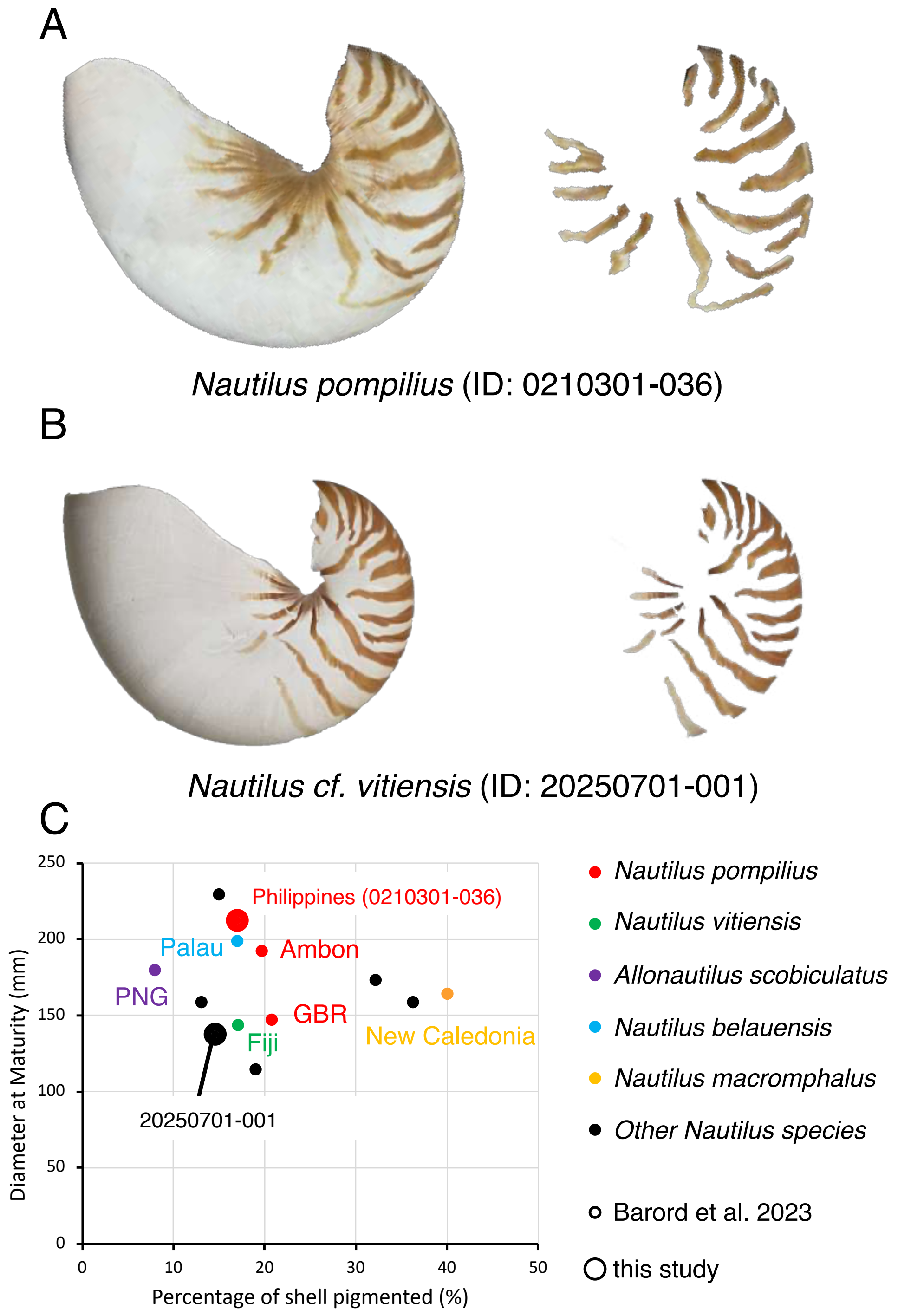

**Fig. S1 Identification of nautilus species based on morphological characteristics.**

Photographs showing the entire shell and pigmented shell regions of two specimens: A) *N. pompilius* and B) *N. cf. vitiensis*. C) Characterization map of *Nautilus* shell morphologies (Barord et al. 2023).

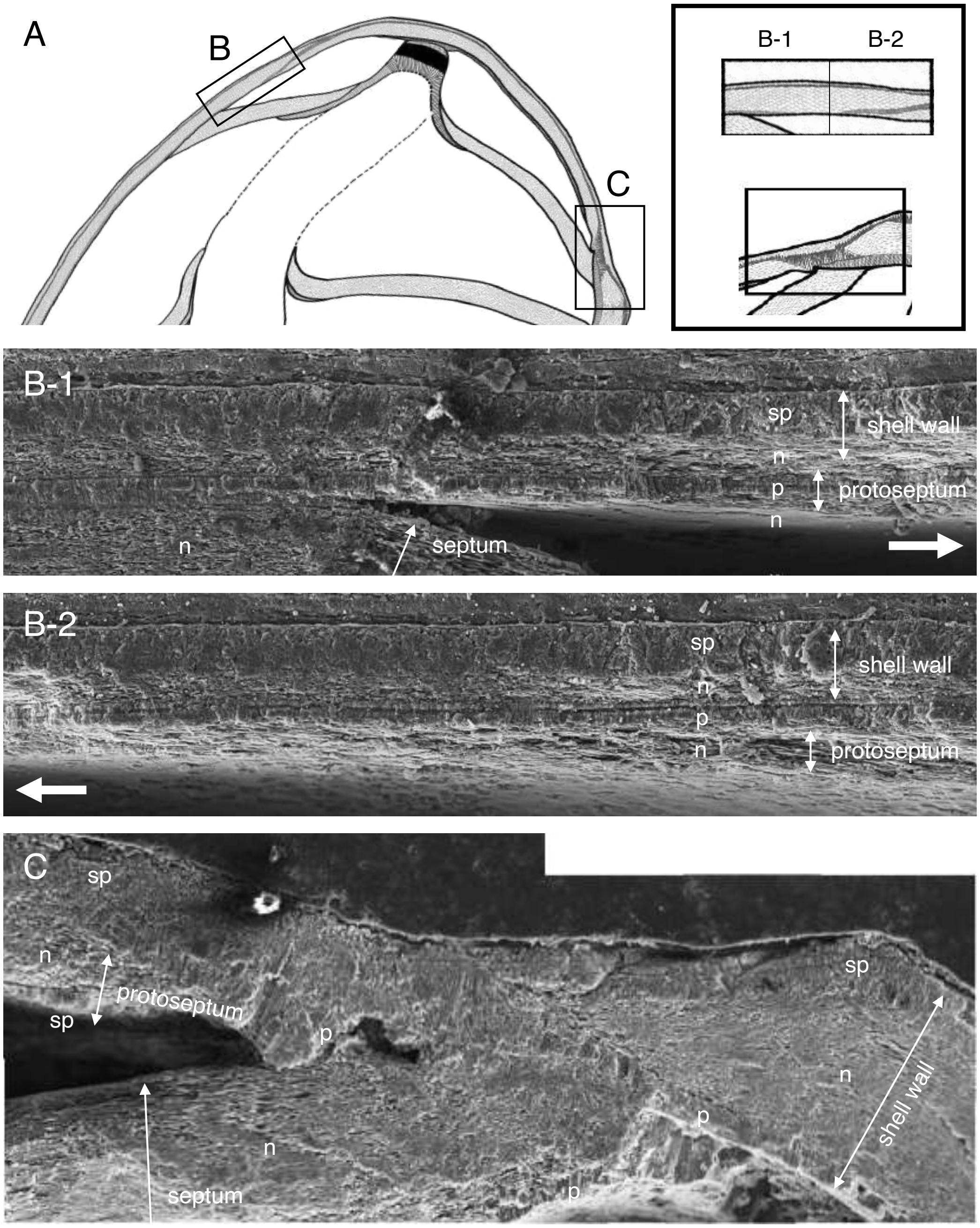
A
B
C
B-1
B-2
B-1
sp
shell wall
n
p
protoseptum
n
septum
n
B-2
sp
shell wall
n
p
n
protoseptum
C
sp
n
protoseptum
sp
p
sp
n
shell wall
p
n
septum
p

8

**Fig. S2. Microstructure of the shell wall during early ontogeny**

Microstructure of the shell wall during early shell development. A) Schematic diagram of the shell at an early developmental stage. SEM images showing microstructures on the 1) ventral and 2) dorsal sides.

Abbreviations: spherulitic and prismatic structure (s-p); prismatic structure (p); nacreous structure (n).
